# Gating modules of the AMPA receptor pore domain revealed by unnatural amino acid mutagenesis

**DOI:** 10.1101/449181

**Authors:** Mette H. Poulsen, Anahita Poshtiban, Viktoria Klippenstein, Valentina Ghisi, Plested Andrew

## Abstract

Ionotropic glutamate receptors (iGluRs) are responsible for fast synaptic transmission throughout the nervous system. Conformational changes of the transmembrane domain (TMD) underlying ion channel activation and desensitization remain poorly understood. Here, we explored the dynamics of the TMD of AMPA-type iGluRs using genetically-encoded unnatural amino acid (UAA) photo-crosslinkers, p-benzoyl-L-phenylalanine (BzF) and p-azido-L-phenylalanine (AzF). We introduced UAAs at sites throughout the TMD of the GluA2 receptor and characterized these mutants in patch-clamp recordings, exposing them to glutamate and UV light. This approach revealed a range of optical effects on the activity of mutant receptors. We found evidence that an interaction between the Pre-M1 and the M4 TMD helix was essential for normal activation and desensitization. Photoactivation at F579AzF, a residue behind the selectivity filter, had extraordinarily broad effects on gating and desensitization. This observation suggests coupling to other parts of the receptor and like in other tetrameric channels, selectivity filter gating.

## Introduction

The AMPA-type glutamate receptor (AMPAR), in common with other ionotropic glutamate receptors (iGluRs), includes glutamate binding domains that connect to a transmembrane ion channel pore. Glutamate binding activates the receptor and opens the channel. Despite the apparent simplicity of this activation principle, the geometries of the receptor involved in different activation states is unclear. Recently two structures from CryoEM of the active AMPAR in complex with the auxiliary subunit stargazin were published (Twomey et al., 2017a) (Chen et al., 2017). However, it remains unclear whether these structures represent fully open channels, or perhaps conformations corresponding to subconductance openings. We currently lack an active state structure for any other subtype of iGluR.

Numerous studies have investigated the structure and function of extracellular domains of AMPARs using pharmacological compounds and/or mutational studies (Sun et al., 2002) (Horning and Mayer, 2004). These studies suggested that desensitization and gating of AMPARs are principally regulated by the conformation of the ligand binding domain (LBD) (Carbone and Plested, 2012), while displacements of the amino terminal domains (ATD) have little effect on AMPAR function (Yelshanskaya et al., 2016a). In contrast to the extracellular domains, comparatively few studies have examined the dynamics of iGluR pore domains. One reason for this deficit is the lack of approaches for capturing movements within the transmembrane domain (TMD).

In tetrameric ion channels, two complementary mechanisms of channel gating have been described. The helical bundle crossing occludes the extracellular end of the AMPAR pore in closed channel structures solved to date (M3 in iGluRs, S6 in potassium channels), which is also tightly occluded in Shaker potassium channels (del Camino and Yellen, 2001) and sodium channels (Oelstrom et al., 2014). But other tetrameric channels including CNG, BK and MthK exhibit either a partially or fully open bundle crossing, with the selectivity filter acting as a principal gate (Contreras et al., 2008; Zhou et al., 2011) (Posson et al., 2013; Thompson and Begenisich, 2012). Even in tetrameric ion channels with minimal architectures such as KcsA, selectivity filter gating underlying C-type inactivation can be coupled to voltage and opening of the bundle crossing, including gating hysteresis (Blunck et al., 2006; Labro et al., 2018; Tilegenova et al., 2017) although see: (Devaraneni et al., 2013). Cysteine accessibility experiments in AMPARs are consistent with a gate at the bundle crossing between resting and open states (Sobolevsky et al., 2004) but only two sites in the M3 helix of AMPARs could be studied with membrane impermeant reagents. Moreover, coupling between the selectivity filter and bundle crossing has not been investigated and may occur in other functional states.

Real-time analysis of receptor activity coupled to chemical crosslinking has the potential to identify sites that have a state-dependent disruption of channel gating. Several techniques to produce crosslinks between parts of the receptor have been used, including disulfide bonding, and the introduction of artificial metal binding sites to bridge subunits (Ahmed et al., 2011) (Lau et al., 2013; Baranovic et al., 2016; Sobolevsky et al., 2004) (Armstrong et al., 2006). However, these approaches require solvent access to the sites of interest, which is not feasible for many sites within the membrane embedded channel domain.

To study the TMD at arbitrary sites, with the aim of mapping the channel gating pathway in an unbiased way, we exploited unnatural amino acids (UAAs). We chose UAAs that are reactive after irradiation with UV light, and that consequently form covalent bonds to nearby protein segments. We used a well-characterised genetic encoding method that has been shown to be highly selective and potent in experiments on 7-TM metabotropic receptors and rhodopsin (Ye et al., 2010; Ye et al., 2009; Ye et al., 2008; Naganathan et al., 2013), potassium channels (Martin et al., 2016; Murray et al., 2016) and NMDA - and AMPA-subtypes of iGluRs (Klippenstein et al., 2014; Klippenstein et al., 2017; Tian and Ye, 2016). We inserted individual TAG (amber) stop codons throughout the TMD of the AMPAR subtype GluA2, rescued these introduced stop codons with p-benzoyl-L-phenylalanine (BzF) and p-azido-L-phenylalanine (AzF) and measured the effects of UV exposures on currents induced by glutamate. We also assessed physical crosslinking with protein biochemistry. These experiments, in concert with analysis of kinetic mechanisms, revealed an unforeseen extent of control of gating and desensitization by both core and peripheral elements of the ion channel domain.

## Results

We hypothesized that structural elements outside the bundle crossing are critically involved in the channel gating reaction. To investigate this hypothesis, we selected 24 sites in the TMD of GluA2 to insert either of two photoactivatable UAAs, AzF or BzF (Figure 1A). The size of AzF is comparable to tyrosine and tryptophan, whereas BzF is bulkier (Figure 1B). Although we preferred to replace aromatic residues, we selected other amino acids at sites that allowed us to cover all four membrane segments, M1-M4. In figure 1 (C & D) we also outline the basic results of the electrophysiological, optical activation and biochemical experiments. This survey already permits some conclusions about the utility and chemistry of UAA crosslinking in the transmembrane segments, as we outline below.

**Figure 1:**
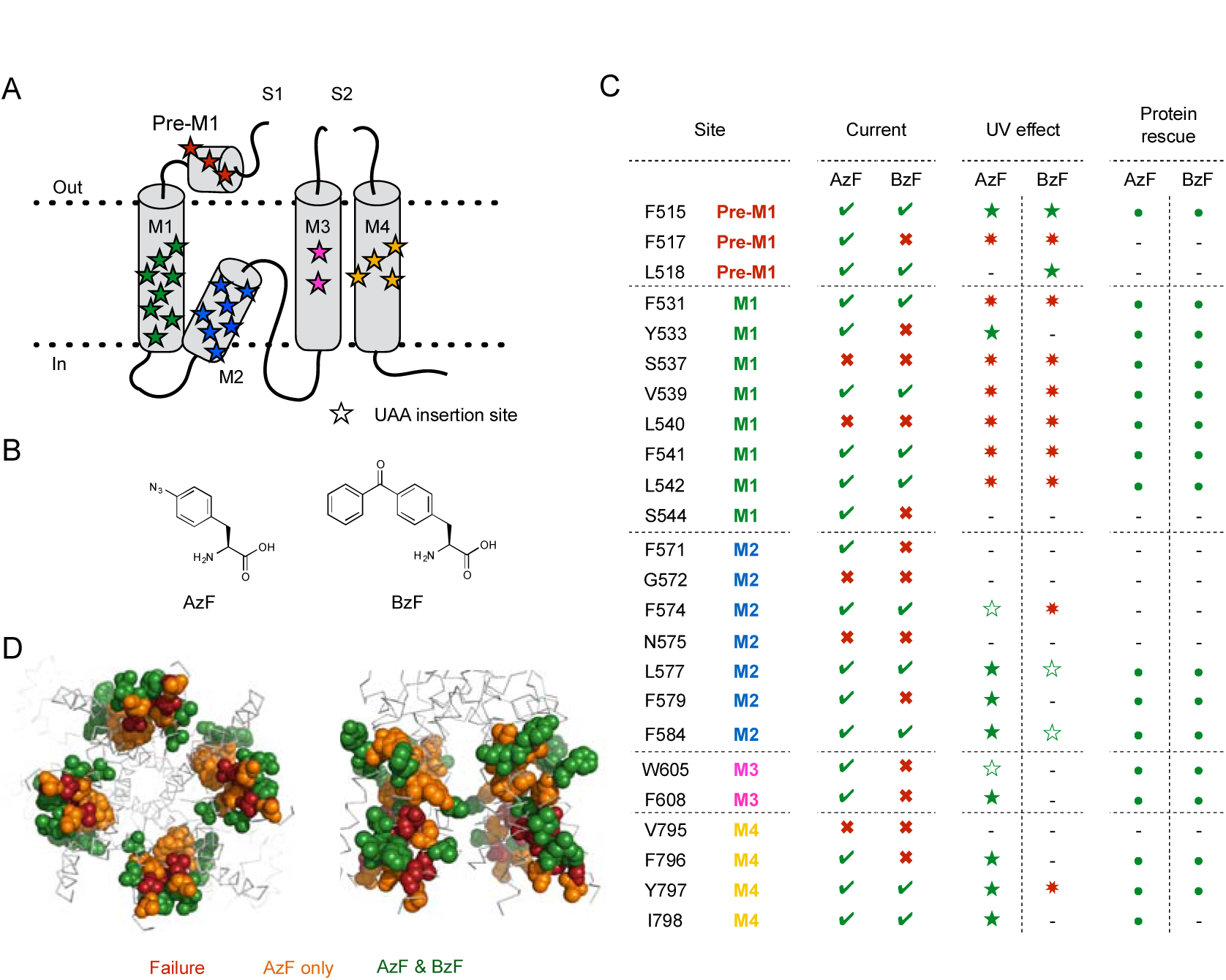
Site-specific incorporation of AzF & BzF. **A** Cartoon of the TMD of one subunit. Stars indicate UAA insertion sites. **B** Chemical structures of AzF and BzF. **C** Summary of electrophysiology, UV effects on currents and expression characteristics. Color-coding of sites according to panel A. Under ‘Current’, green ticks indicate constructs for which glutamate induced currents could be detected, and red crosses indicate constructs where no currents could be recorded for AzF or BzF incorporation. Receptors were exposed to UV light in the resting and/or desensitized state during electrophysiological recordings. Green filled stars indicate a specific UV-triggered effect and red asterisks indicate no apparent effect of UV light. Unfilled green stars indicate constructs (GluA2-F574AzF, - W605AzF, - L577BzF and - F584BzF) where large rundown in the current amplitude precluded quantitation of any possible slow concurrent UV effects. Recordings of the GluA2-I798BzF mutant had rapid rundown and leak currents that were larger than usual. The currents for GluA2-F571AzF and - S544AzF mutants were too small to permit analysis of UV-driven effects (indicated by hyphens). Western blot analysis indicated that “rescue” of translation was successful for all the constructs tested, indicated by filled green circles. A hyphen also indicates no test. **D** Representation of insertion sites shown as color-coded spheres in a GluA2 crystal structure (PDB ID: 3kg2). Red spheres indicate failure of both AzF and BzF to rescue functional receptors (‘Current’ column in panel C), whereas orange spheres indicate successful insertion of only AzF and green spheres indicate successful insertion of both AzF and BzF.

First, we found insertion sites in all helices at which we could rescue functional channels at the plasma membrane. At 19 of the 24 sites, typical fast glutamate-activated currents could be recorded from channels harbouring AzF. At 11 of these sites, BzF also produced functional channels. There were no sites at which the bulkier BzF could preferentially rescue channel function. Viewed on the basis of crystal structures of GluA2, a further aspect of functional rescue was apparent. Sites that readily accommodated both amino acids tended to be found at the circumference of the membrane domain, whereas sites that produced non-functional channels for both AzF and BzF tended to be centrally located around the channel pore. An intermediate layer of sites was permissive for AzF alone (Figure 1D).

At 9 of the 19 sites rescued by AzF, we could detect a robust alteration of channel gating upon exposure to UV light. We explore in more detail the nature of these changes below. Strikingly, we could not determine a robust effect of BzF photoactivation on currents for any of the sites tested, except for the peripheral sites at F515 and L518 in the “Pre-M1” helix, which are likely to be outside the plasma membrane in any case. We did detect a weak effect of UV exposure when BzF was incorporated in the M2 helix at position 577, however, the effect was hard to separate from current rundown. The absence of distinct UV-induced changes on receptor function for constructs containing BzF is surprising in light of experiments in which membrane domains were crosslinked with Benzophenone [PMID: 11524017], but may relate to water accessibility in such sites. The only water accessible sites in the AMPAR pore domain are likely to be in the M3 helix (Sobolevsky et al., 2005) (Sobolevsky et al., 2003) (Salussolia et al., 2011) and BzF failed to produce functional channels at these sites.

A possible confounder of these results was that a lack of glutamate-activated current expression was because of a failure to translate the polypeptide chain. Therefore, we performed biochemical experiments to assess rescue of expression. These experiments confirmed that for both AzF and BzF, inclusion of the UAAs and the requisite synthetase and exogenous tRNA were sufficient to strongly enrich expression of the full-length subunits (Figure 1C, Supplementary Figure 1 and Supplementary Table 1, range 5- to 3500-fold increase in band intensity for 10 mutants). On averag e, wild-type receptors showed no change in expression level in the presence of the UAAs and the incorporation machinery. These experiments do not provide information about the maturity of the tetrameric form of the receptor, or about surface expression, but do indicate that deficits due to UAAs were either in assembly and/or gating, not in a gross absence of translation of the full-length polypeptide chain.

To assess whether functional receptors rescued by AzF and BzF were valid congeners of wild-type receptors, we assessed their desensitization, deactivation and recovery from desensitization. Example traces are plotted in Figure 2 and analysis of the kinetics of nine mutants is provided in Supplementary Table 2. Out of all the mutants tested, only insertion of AzF at position 798 had any appreciable effect on kinetics slowing the rate of entry to desensitization to 50 ± 5 s^−1^ (*n* = 6 patches) compared to 120 ± 10 s^−1^ for wild-type receptors (*n* = 36 patches). Overall, the range of deactivation rates was from 1200 s ^−1^ to 2200 s^−1^, compared to the average wild-type value of 1600 ± 120 s^−1^ (*n* = 27) and the recovery from desensitization ranged from 45 s ^−1^ to 70 s ^−1^ compared to the average wild-type value of 55 ± 5 s^−1^ (*n* = 17) (Figure 2; Supplementary Table 2). Therefore, we took the mutants in their basal state, before any UV exposure, to have gating largely representative of GluA2 wild-type (WT) channels.

### Patterns of UV-induced inhibition

Previously, we generated photo-inactivatable AMPARs by inserting BzF in the extracellular domains of GluA2 (Klippenstein et al., 2014). Due to their incorporation sites, these mutations were expected to trap an inactive state if they formed crosslinks. Here, we chose sites on a pseudo-random basis, without any particular expected photo-crosslinking effect. For five of the AzF mutants, the effect of UV illumination was a rapid, irreversible loss of the peak current response (Figure 3A-D; Supplementary Table 3). Independent of the location (in any of the four membrane segments, M1-4), we could inhibit the peak current by up to 95%. The fastest inhibitory action was observed for the F608AzF mutant, with a time constant of 1.5 s for cumulative exposures to UV (in intervals of 200 ms per episode) for reduction of the peak current to 4 ± 1% of its original value (barely distinguishable from background noise; *n* = 17,Figure 3C and Supplementary Table 3). We were able to control the speed of inactivation by changing the intensity of the UV light (50% to 100%) or the time interval of UV exposure per episode (50 ms and 200 ms) (Supplementary Figure 2). We exposed receptors to UV light in resting and desensitized states by opening the shutter at the appropriate stages of each episode. To examine the active state, we initially blocked desensitization with cyclothiazide (CTZ), but found that CTZ itself could induce a UV-sensitive inhibition (Supplementary Figure 3) that varied from batch to batch. To avoid this problem, we instead performed experiments on the background of the L483Y mutation for all constructs, with the exception of F608AzF-L483Y, which did not express. For this particular construct, F608AzF, the UV induced inhibition in presence of CTZ was much faster than the CTZ-driven UV dependent inhibition of GluA2 WT (see Supplementary Table 3). Surprisingly, the effects of UV exposure were independent of the functional state of the receptor. Values for all the constructs tested that showed UV dependent inhibition are reported in Supplementary Table 3. As previously reported (Klippenstein et al., 2014), we controlled for non-specific rundown of currents by pausing the UV exposures in the course of some experiments, observing that the peak current remained stable, and then reverting to UV-driven inhibition.

**Figure 2.**
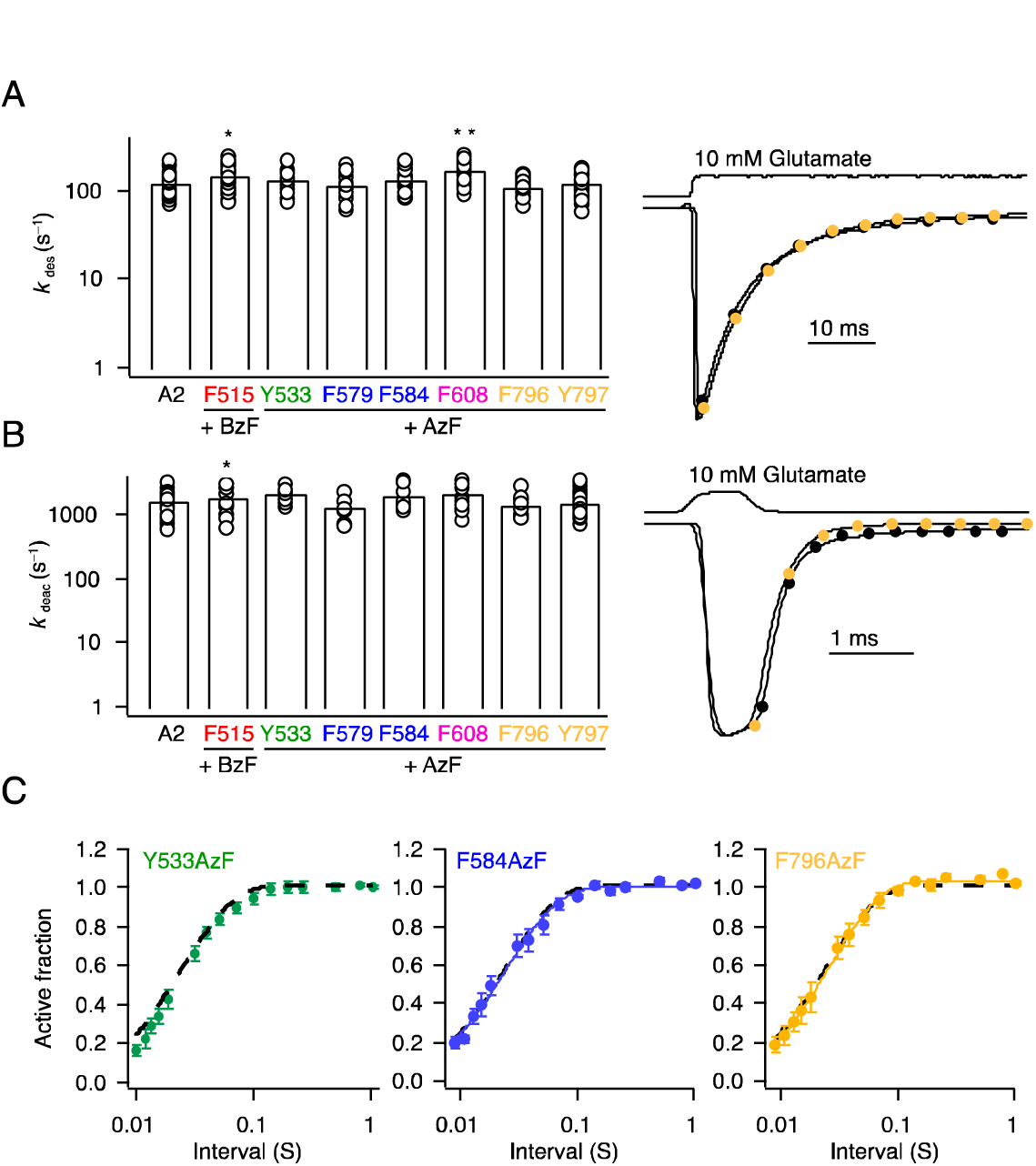
Kinetics of GluA2 receptors harboring AzF or BzF in the transmembrane domain. **A** Bar graph (*left*) summarizing desensitization rates of selected GluA2 constructs with AzF or BzF. ^**^Significant difference GluA2 WT vs. mutant (p < 0.001, t-test), ^*^ Significant difference GluA2 WT vs. mutant (p < 0.05, t-test). Traces (*right*) illustrating the rate of desensitization of GluA2 WT (*black*) and F796AzF (*yellow*). **B** Bar graph (*left*) summarizing deactivation rates for selected constructs with AzF or BzF after a brief (1 ms) pulse of 10 mM glutamate. Traces (*right*) illustrate the rate of deactivation of GluA2 WT (*black*) and F796AzF (*yellow*). **C** Pooled data for recovery from desensitization of GluA2 WT (black dotted line), GluA2-Y533AzF (green), - F584AzF (blue) and - F796AzF (yellow). See Supplementary Table 2 for a summary of rates.

**Figure 3.**
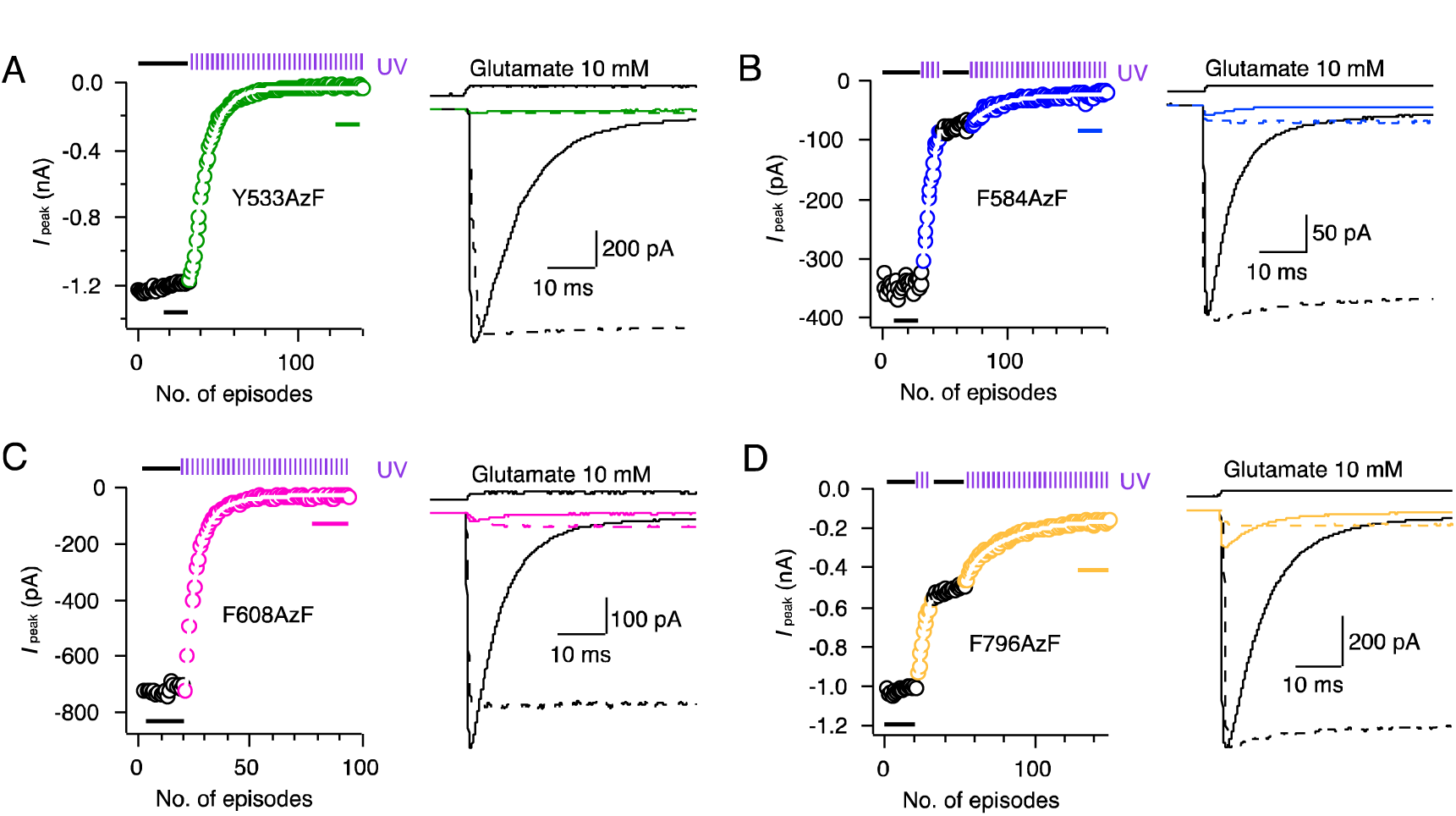
UV-triggered inhibition of glutamate-induced currents. **A-D** *Left*; Examples of kymograms illustrating the time course of receptor photo-inactivation for the selected GluA2 constructs. Each episode included a 400 ms application of 10 mM glutamate (each circle represents the peak current of the response). A 200 ms exposure of UV epi-illumination was made in each episode (indicated in the kymogram by violet pulse trains and colored circles). The rate of peak current reduction was monoexponential (white outlined fits). There was no significant difference between monoexponential half-times in different states of activation. *Right*; Traces representing averages of 5-20 responses to glutamate before UV exposure (black trace) and after UV exposure (colored) in either the resting or desensitized state (solid lines, taken from points indicated by bars in kymogram) and fully active state (corresponding L483Y mutant, dotted line). It was not possible to record any glutamate induced current from GluA2-L483Y-F608AzF, therefore CTZ was used to block desensitization for this construct.

Such a consistent pattern of inhibitory action might argue for a non-specific loss of channel function, but several observations counter this proposition. As shown previously (Klippenstein et al., 2014), GluA2 WT receptors were insensitive to UV light (Supplementary Figure 4A), as were some mutants. For example, F531AzF was reliably rescued by AzF but showed no UV sensitivity at all (Supplementary Figure 4B), speaking against non-specific photo-destructive effects based on AzF. A further control was provided by the F584TAG mutant in the absence of any UAA. All permissive AzF mutants had some degree of readthrough (that is, background rescue in the absence of UAA). The F584TAG mutation had the largest readthrough currents on average, similar in magnitude to those in the presence of AzF (Supplementary Table 4). However, unlike responses from receptors containing the AzF residue, the responses obtained (in the absence of AzF) were entirely insensitive to UV illumination. A typical recording from the F584TAG mutant from a cell cultured in the absence of AzF is shown in supplementary figure 4C.

Although these controls gave us confidence that the inhibitory effects were site-specific, the behaviour of several other mutants provided more compelling evidence. For example, the F796AzF mutation in M4 was inhibited by UV, but inhibition was incomplete reaching only about 85%. In addition, we found that incorporating AzF in M2 at position F579 also resulted in inhibition upon UV illumination, but only to 75% and with a concomitant increase in steady-state current (see below and Supplementary Table 3). This suggests that the effects of AzF activation within the transmembrane domain are site-specific.

### Photopotentiation

As would be expected from an unbiased screen of the gating region, in addition to inhibition, we also found four mutants that had a strong potentiating effect on function. For F515BzF the peak current was on average increased 1.6-fold, with a larger 2.5-fold increase in the relative steady-state current compared to the peak current size (Figure 4 and Supplementary Table 3). Peak current was also increased for the non-desensitizing LY mutant indicating an effect on gating, not a simple block of desensitization (Sun et al., 2002). Glutamate-activated peak and steady-state currents were similarly increased for L518BzF by 1.3-fold and 7.8-fold, respectively (Figure 4). As expected from a stabilisation of the open state, paired measurements in the same patch (before and after UV exposure) showed the rate of entry to desensitization

**Figure 4.**
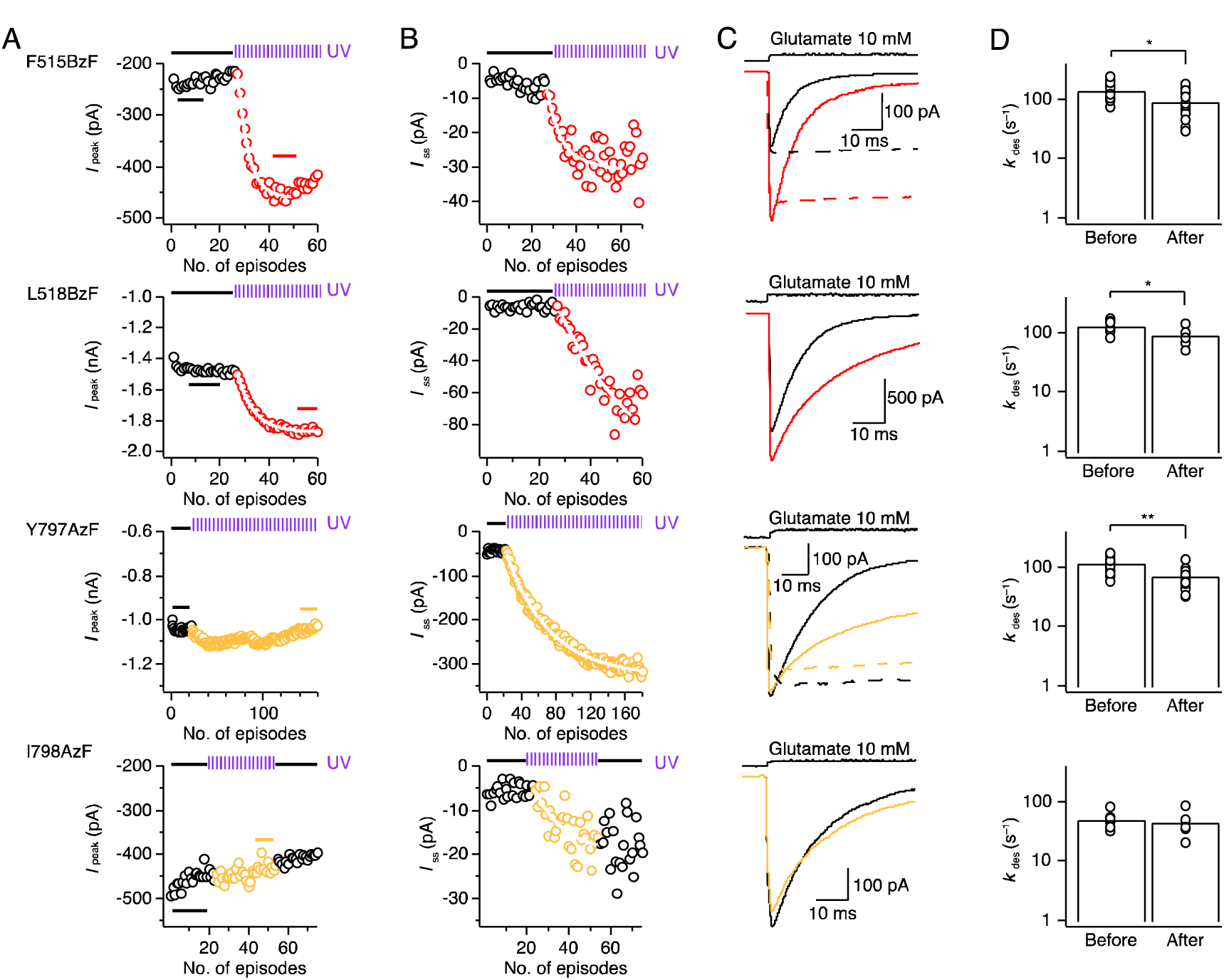
UV-triggered potentiation of AMPAR responses. **A-B** Example kymograms illustrating the time course of the potentiation of peak (**A**) and steady state (**B**) current for GluA2-F515BzF (*top row*), GluA2-L518BzF (*second row*) GluA2-Y797AzF (*third row*) and GluA2-I798AzF (*bottom row*). The rate of peak current potentiation was monoexponential (white outlined fits). **C** Example current traces representing averages of 5-20 responses to glutamate before UV exposure (black trace) and after UV exposure (colored) in either the resting or desensitized state (solid lines, taken from points indicated by bars in kymogram). Representative currents from the corresponding L483Y mutants (active state UV exposure, dotted lines) are shown for F515BzF and Y797AzF. D Bar graphs summarize desensitization rates in 10 mM glutamate of GluA2-F515BzF, - L518BzF, - Y797AzF and - I798AzF before and after UV exposures (see Supplementary Table 2 for a summary of rates). was slowed (from 140 ± 10 s^−1^ to 90 ± 10 s^−1^, *n* = 19 and 130 ± 10 s^−1^ to 90 ± 10 s^−1^, *n* = 7, for F515BzF and L518BzF, respectively). An analysis including both paired and non-paired data is in Supplementary Table 2.

A similar effect on receptor activation behaviour was seen for AzF introduced in M4 at position Y797, with an increase in the steady-state current (3.7-fold), presumably corresponding to the block of desensitization (from 120 ± 5 s^−1^ to 70 ± 5 s^−1^, *n* = 24) (Figure 4). Similarly, the relative steady-state current increased, albeit more variably, following UV exposure when AzF was incorporated into the neighbouring M4 position at I798 (fold increase of 7 ± 3, Figure 4 and Supplementary Table 3). The extent of photopotentiation of the steady-state currents did not correlate with the initial size of the relative steady-state currents before the application of UV light (*R*^2^ range from 0.06 to 0.2, Supplementary Figure 5). Notably, desensitization was slowed by incorporating AzF at position I798 and did not change with exposure to UV light (from 50 ± 5 s_−1_ to 45 ± 10 s^−1^, *n* = 6) (Figure 4). Since the I798AzF construct showed pronounced rundown, we were unable to determine deactivation and recovery from desensitization kinetics before and after applying UV light.

### The interaction between the Pre-M1 and M4 helices

The confluence of the functional results in terms of potentiation of constructs with BzF in Pre-M1 (F515BzF, L518BzF) and AzF in M4 (Y797AzF, I798AzF) led us to ask what structural dynamics could produce these effects. Inserting the mutated residues into the closed state structure of GluA2 provides a striking hypothesis: reciprocal crosslinking between these two sites is physically plausible (Figure 5A). The BzF substitution at position 515 is located on the outer face of the Pre-M1 helix, but a rotation or outward bloom of this “collar” (Twomey et al., 2017a) could allow the photoactivated BzF radical to contact the M4 helix. Likewise, the AzF at 797 is likely buried within the membrane but conformational change could allow it to reach multiple crosslinking partners. Rotations of M4 could permit crosslinking onto M3 or M1 of the neighbouring subunit, but approach of the Pre-M1 helix from the same subunit, as envisaged above, could also bring this residue into potential contact.

Intersubunit crosslinks, like those previously seen for the S729BzF mutant (Klippenstein et al., 2014) were negligible for GluA2 WT receptors and all mutants that we tested (Supplementary Table 1). To test our hypothesis of reciprocal crosslinking between Pre-M1 and M4 within subunits, we used a principle based on a previously published study (Xu et al., 2013), as illustrated in Figure 5B. Using antibodies to label both N - and C - termini, we used quantitative Western blotting to determine the protection against protease digestion afforded by crosslinking. We expected to detect intrasubunit crosslinking if the UV-activated UAA was able to physically connect fragments divided by an inline tobacco-etch virus (TEV) cleavage site. Following brief UV exposures of 2 and 5 minutes, the F515BzF mutant harbouring a TEV cleavage site showed a small but reproducible increase in the protected monomer fraction (Figure 5C-D). Based on the location of F515 in structures of GluA2, and our failure to detect any increase in dimers or hig her order olig omers, this result indicates cross-linking to M4 of the same subunit. The Y797AzF and I798AzF mutants did not show detectable monomer protection (Figure 5C-D), indicating that the effects in electrophysiological experiments likely arise either from a ring-expansion of the phenyl ring, or crosslinking to lipid. Both interpretations require a movement of M4 (see discussion). These exposures to UV light were about 100-fold less intense than those experienced by receptors during patch clamp experiments and epi-illumination by UV (Klippenstein et al., 2014), therefore the total exposure over a few minutes should be similar to those in electrophysiology experiments. Longer exposures of 15-30 minutes, which we previously used to test for dimer (and higher order) fractions, were not relevant for the rapid changes in the gating properties that we were concerned with.

**Figure 5.**
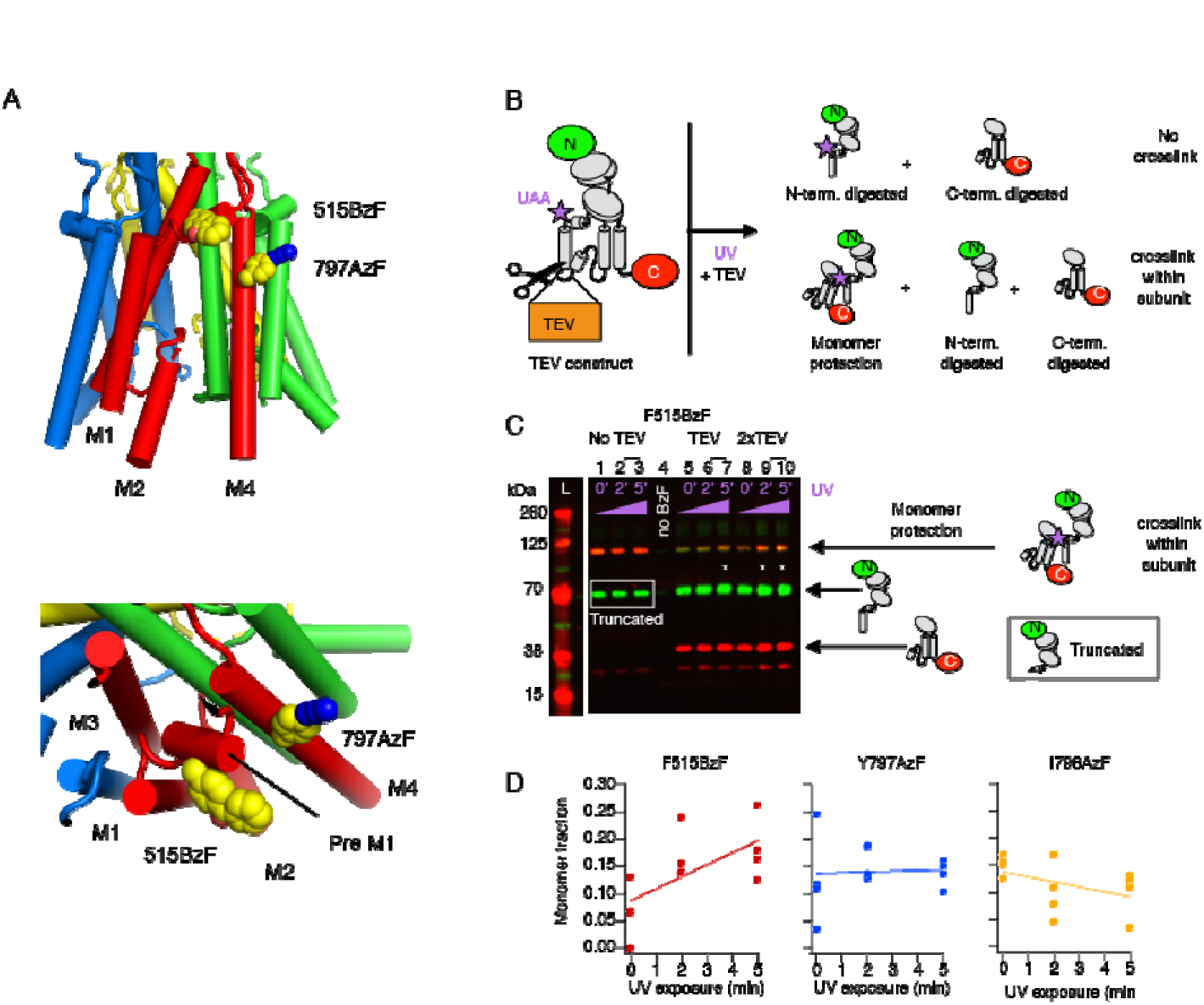
Intra-subunit crosslinking by BzF. **A** Structure of the TMD with AzF and BzF incorporated at sites 797 and 515, respectively, illustrating the close proximity of the two sites. **B** Cartoon of the TEV construct and fragments generated by TEV protease treatment. A site for TEV protease recognition was introduced in the M1-M2 intracellular loop of the GluA2 subunit as well as a C-terminal FLAG-tag epitope and a TAG mutation for incorporation of UAA (*violet star*). Covalent bridging within a subunit of the two fragments arising from TEV protease treatment should “protect” a monomeric subunit band on western blots. **C** Exemplary western blot showing monomer protection of GluA2-F515BzF. For all conditions the band at 63 kDa corresponds to GluA2 truncated at residue 515. The truncated band is undistinguishable from the digested N-terminus when TEV protease is added (64 kDa). *Lane 1-3*: Quantitation of the rescue of F515TAG construct in cells by BzF, showed a monomeric band at 100 kDa. A band from subunits truncated at the TAG site (63 kDa) is present, presumably pulled down in FLAG purification with full-length subunits. *Lane 4*: The omission of only the UAA results in no rescue of monomeric band. A band corresponding to subunits truncated at the TAG site can be visualized on the blot, however the band is very faint, presumably due to a lack of any FLAG epitope. Lane 5-7: Quantitation of GluA2-F515BzF treated with TEV protease showed an increase of monomer fraction with longer exposure to UV. Lane 8-10: Exposing GluA2-F515BzF to twice the amount of TEV protease (relative to lane 5-6) led to an increase in protection of monomers, indicating more crosslinking events. D Summary of the monomeric fraction plotted against the UV exposure time. Only insertion of BzF in position 515 showed an increase in monomer protection over time.

### Structural mapping of photoactive sites

The effects of UV exposure on the peak, relative and absolute steady-state current of rescued receptors are summarised as bar plots in figure 6. We mapped these effects onto a structure of the GluA2 TMD (PDB: 3KG2; Figure 6B). These plots suggested gradients in the peak and steady-state current effects, but any correlations are weak (*R*^2^ = 0.3 for the fold change in peak current against radial distance; Supplementary Figure 6). The weakness of these relations is not surprising, given that the chemistry of individual side chain environments as well as local motions must influence the effect of a given residue on channel gating motions.

Structures of GluA2 in resting, active and desensitized states are now available, so we next asked if the displacement of residues between structures could be related to the photo-activated effects on currents. We selected two nominally resting structures bound with antagonists (5L1B and 5VHZ), two active state structures with open pores (5VOT and 5WEO) and two desensitized state structures (5VOV and 5WEK). These structures were obtained in both the presence and absence of different auxiliary proteins, which may contribute to variations between them. We aligned them on the basis of their membrane embedded segments and measured per-subunit distances between residues in structures, and also took radial displacements by halving distances between diametrically-opposed subunits (e.g. A and C, and B and D; Supplementary Figure 7).

Measurements of distances between residues in different states produced a complex picture indicating considerable structural plasticity both within and between functional states. The two open state structures are quite similar, but the variability between the closed state structures was high (Supplementary Figure 7C) and accounts for almost all the variability found in C-alpha positions between “closed” and “open”. Notably, the most variable regions are the selectivity filter and the top of the M4 helix.

Finding no clear relation between geometry and the functional effect, we produced a 2-D map of gating changes after UV exposure (Figure 7A), plotting the peak current change against the relative steady-state current change. The plot delineates a cluster of null mutants, and two clear groups for which functional changes emerged, whereas the F579AzF mutant was a striking outlier. In the first of the two groups, peak current was inhibited and the steady-state current concomitantly increased (Figure 7A, purple circles). In the second group, a robust increase in steady-state current was accompanied with either increase or no change in the peak current (Figure 7A, green circles). Notably, F579AzF could be described as an outlier in plots of peak current and steady-state current against geometry (Supplementary Figure 6).

Simulations based on a simple single binding site model of AMPARs (Carbone and Plested, 2012; Carbone and Plested, 2016) can mimic the behaviour of these two groups (Figure 7B-D). The behaviour of the first group of mutants on the 2-D plot was reproduced by progressively reducing the lifetime of the desensitized state AD2 or by allowing the desensitized state to become weakly conductive (Figure 7B-D). The kinetic behaviour of the second group of mutants was reproduced by altering the channel shutting rate α or the opening rate β (Figure 7B-D). The quantitative agreement between the 5-fold increase in the steady-state current accompanied by a 50-fold reduction in the peak current predicted by this model, and the effects on actual mutants (e.g. 608AzF) is notable. Most importantly, these simulations indicated that crosslinking at the F579 position is complex in nature, and must involve multiple effects on kinetic model parameters, where both the lifetime of desensitization and channel opening rate change (Figure 7B-D). A further manipulation, allowing the desensitised state to become progressively weakly conductive gave the best description (Figure 7B-D).

Figure 7E (and Movie 1) illustrates four classes of residues that we could segregate by analogy to kinetic models of AMPAR activation. The residues form contiguous clusters in the TMD, suggesting that they correspond to functional modules that execute distinct gating functions. Such modules strongly resemble the “conformational wave” model of AChR receptor gating (Grosman et al., 2000). We do not know the extent to which these “modules” are independent, or coupled within a common pathway, or the extent to which they are disrupted or highlighted by crosslinking.

**Figure 6.**
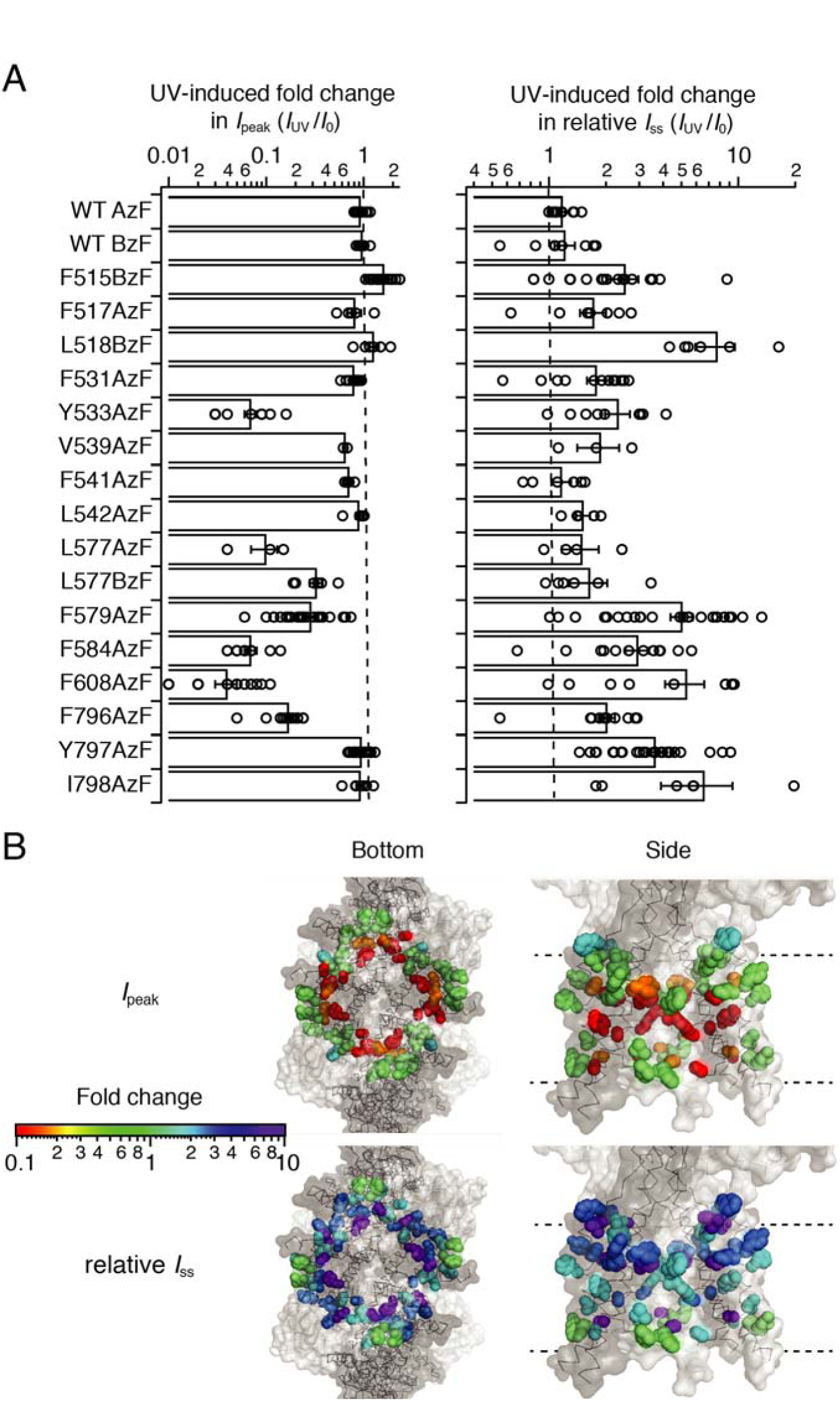
Segregation of sites by UV effect. **A** Bar graph (*left*) representing the summary of the change in peak current before (*I*_0_) and after UV (*I*_UV_) of selected GluA2 constructs with AzF or BzF incorporated in the TMD. Small reductions in peak current (like for GluA2-F517AzF) are likely due to rundown and not related to the application of UV. Bar graph (*right*) summarizing the fold change in steady-state current relative to the peak current. **B** Bottom and side view of structures showing AzF and BzF insertion sites as spheres colored according to their UV-dependent changes in peak current amplitude (*upper*) and relative steady-state current (*lower*). Each site is highlighted in color in all four subunits. The color scale (*left*) is representing the colors used to show the fold change measured for each construct.

**Figure 7.**
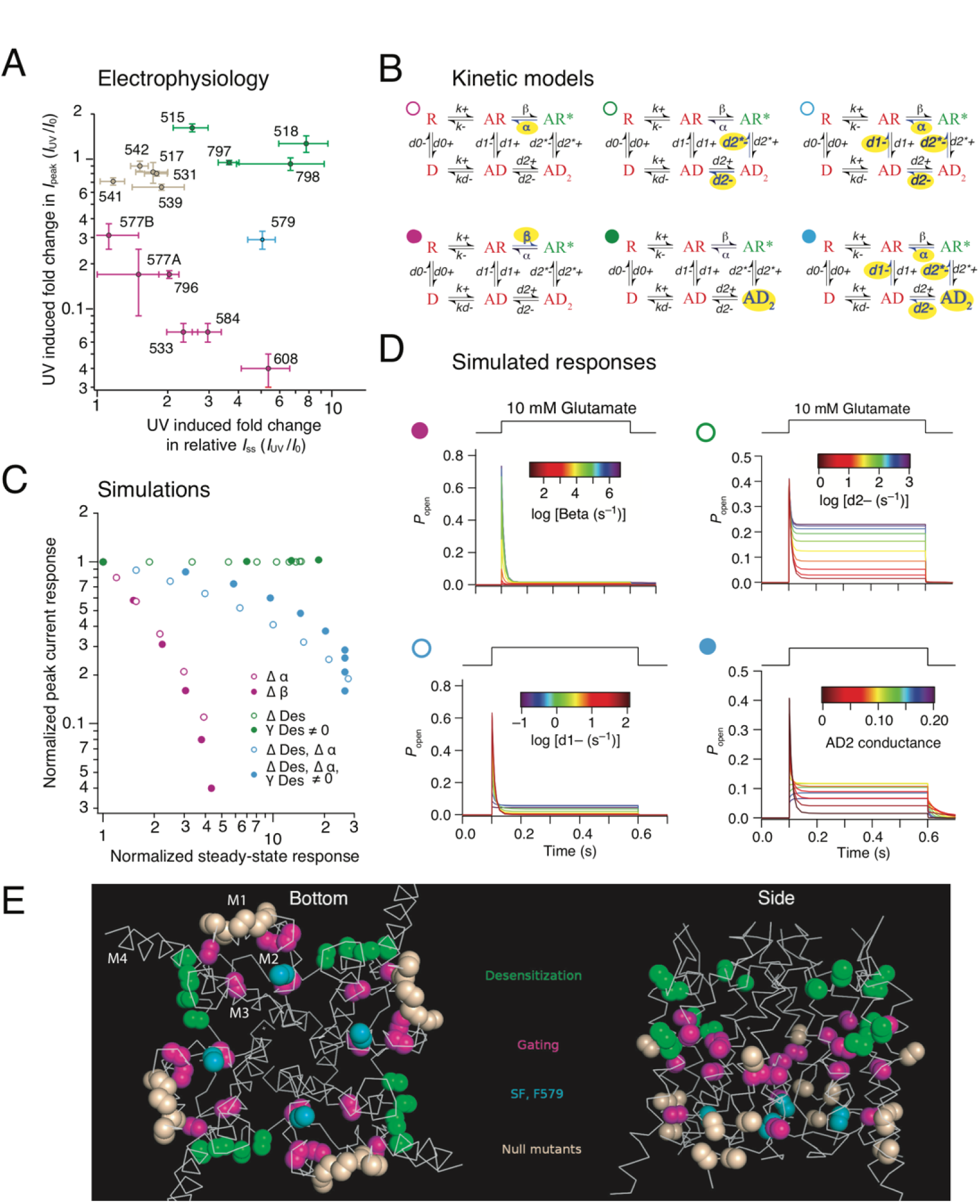
Segregation based on gating properties. **A** 2-D plot shows the relation between the mean UV-induced changes in peak and relative steady-state current before (*I*_0_) and after UV (*I*_UV_) at each site. Sites are color-coded according to the effect: null in wheat; potentiation in green; inhibition in purple and intermediate (579) in cyan, the same color code applies to each panel. 577A and 577B denotes the insertion of AzF and BzF, respectively, at this particular site. **B** Simplified single binding site models of AMPAR gating. Open state (AR^*^) is green, shut states (including desensitized states D, AD and AD_2_) are red. Rates or states that were varied in each simulation are highlighted in yellow. **C** A 2-D plot, resembling the 2-D plot in panel A, but derived from simulations of the kinetic models described in panel B. Progressive alteration of the channel shutting and opening rate (α and β, purple circles), changes in desensitization rates (d2, open green circles) or AD2 becoming conductive (filled green circles). Changes to both gating and desensitization are needed to obtain the intermediate behavior (cyan circles). **D** Simulated responses from kinetic models (indicated with colored circles as in panel B) used to construct panel C, with traces colored according to the rate constant indicated. **E** Gating modules in the AMPAR pore. The major classes of mutants form contiguous modules; a desensitization module (green; collar), a gating module (magenta; bundle crossing) and peripheral mutations with no effect (null mutants, wheat). The selectivity filter mutant with complex behavior is F579AzF (cyan).

### Complex role in receptor kinetics of the F579 site

A complicated kinetic model was needed to mimic the position of F579AzF in the 2-D gating plot (Figure 7). Therefore, we reasoned that complex UV-driven changes in gating at this site might be detectable in patch clamp recordings. To address this point, we pursued a more detailed investigation of kinetics, including recovery from desensitization, at both an intermediate time point in cumulative UV exposure (~2 s) and after saturating exposure (8s).

Kymograms of the UV effects on peak current and steady-state response are shown in Figure 8A. As in our initial analysis, peak current inhibition by UV was incomplete, reaching only about 75% (Figure 8A and Supplementary Table 3). This was also true for the non-desensitizing L483Y variant, indicating that reduced channel gating, rather than increased desensitization, was responsible for this inhibition. In these experiments, the rate of entry to desensitization was reduced by almost one third after saturating UV exposure time of 8s (from *k*_des_ = 130 ± 15 s^−1^ to *k*_des_ = 45 ± 10 s^−1^, *n* = 9, Figure 8B), whereas intermediate cumulative exposure time of 2s showed desensitization rates similar to control (*k*_des_ = 110 ± 10 s^−1^). Paradoxically, steady-state currents activated by 10 mM glutamate increased to about 10% of the peak (Figure 8B). Likewise, the deactivation rate following a 1 ms pulse of 10 mM glutamate was also substantially slowed (from 1410 ± 140 s^−1^ to 700 ± 140 s^−1^ (2s UV) and 500 ± 140 s^−1^ (8s UV), *n* = 9, Figure 8C). Both these effects on gating developed strongly at early stages of UV exposure (that is, after only 2 s).

As expected from our kinetic modelling, the effect on desensitization was not simple. To this end, also recovery from desensitization was slowed (from *k*_rec_ = 40 ± 5 s^−1^ to *k*_rec_ = 20 ± 5 s^−1^(2s UV) and *k*_rec_ = 15 ± 5 s^−1^ (8s UV), *n* = 9, Figure 9A-B). One caveat for this measurement is that control measurements on the GluA2 WT receptor, expressed on the background of AzF and its cognate synthetase, showed a small slowing of recovery after UV exposure (from 55 ± 4 s^−1^to 38 ± 2 s^−1^, *p* = 0.008, paired t-test, n = 9 patches, Supplementary Table 2). The effect on F579AzF mutants was much more substantial, and quite different to control measurements, with a “quiescent phase” where no effective response was detected for about 30 ms following the conditioning pulse (Figure 9A). Measuring recovery was hampered by the peak current inhibition but also by the development of an unusual long decay in the current after the desensitizing pulse. This “off-relaxation” following desensitization was much slower after only a brief UV exposure, (from *k*_off_ = 150 ± 60 s^−1^ to *k*_off_ = 30 ± 5 s^−1^ (2s UV) and *k*_off_ = 35 ± 5 s^−1^ (8s UV), *n* = 9, Figure 9C). These measurements also revealed that the steady-state current increased in absolute magnitude. There was no concomitant effect on the resting state conductance - the patches did not become leaky.

By comparing effects on kinetics at intermediate and saturating UV exposures, it was clear that the onset of UV-driven effects in F579AzF was not coherent. The slowing effect on the tail current (*k*_off_) and deactivation as well as the slowing of recovery from desensitization were pronounced already at 2 s, whereas effects on the steady-state current and the rate of entry to desensitization developed later (Figure 10A). These distinct time courses are further evidence that photoactivation of the F579AzF mutant has an unprecedent effect on multiple functional states of the AMPAR. This observation is consistent with a high structural variability at the selectivity filter region (at the cytoplasmic end of the channel; Figure 10B and C) both between and within states.

**Figure 8.**
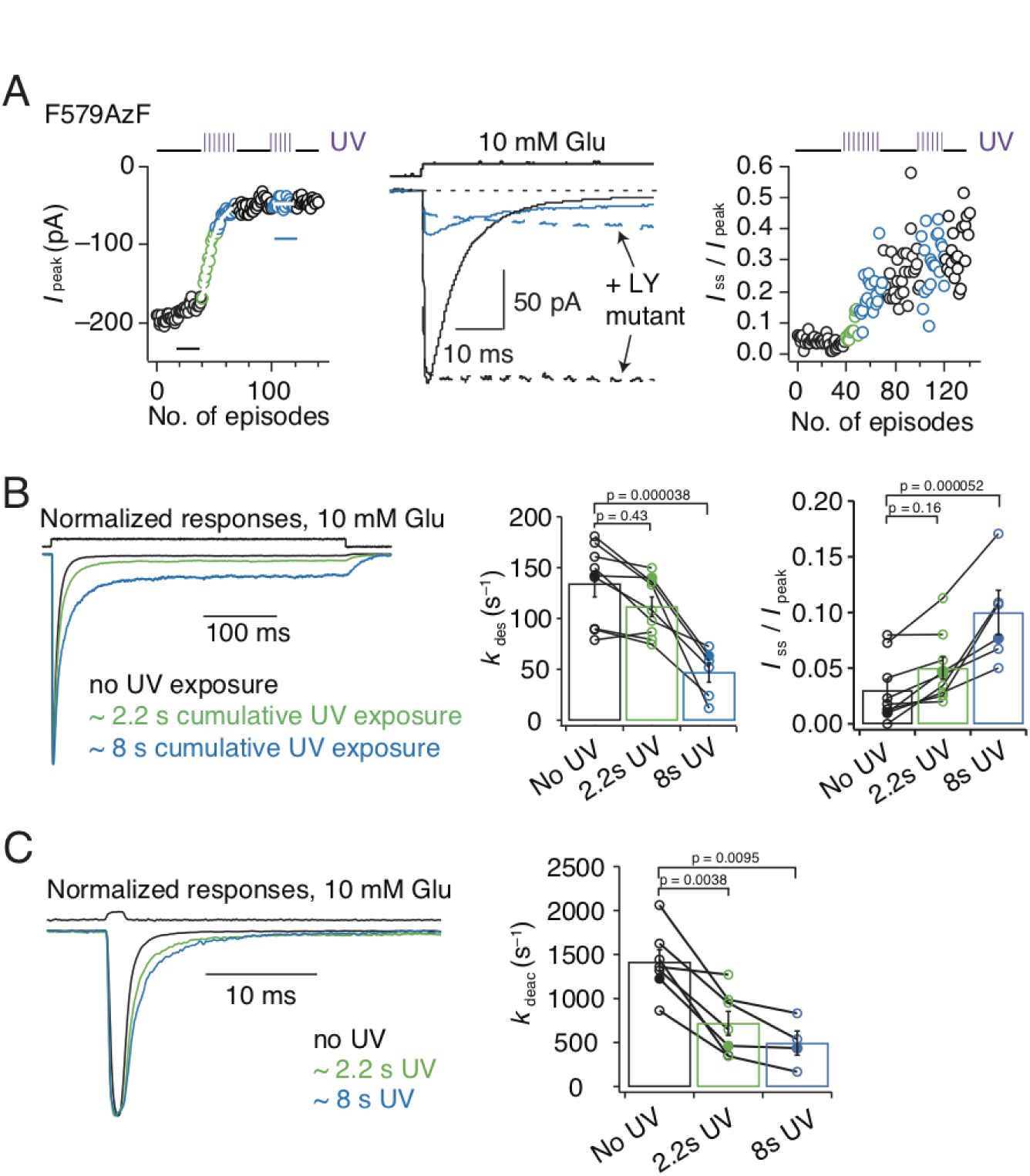
Broad role of the M2 segment in activation and desensitization. **A** Kymograms illustrating the time course of incomplete inhibition for GluA2-F579AzF peak current responses to 10 mM glutamate (*left*) and increase in the relative steady-state current (*right*). Example current traces (*middle*) representing averages of 10 responses to glutamate before UV exposure (black trace) and after UV exposure (cyan) in desensitized state (solid lines) and fully active state (corresponding L483Y mutant, dotted line). **B** Exemplary normalized traces for desensitizing responses to a 400 ms glutamate pulse, before (black), during (2.2 s, green) and after (8 s, blue) UV exposure. Bar graph shows rates of desensitization, relative steady-state current (*I*_ss_/ *I*_peak_) ± SEM before, after 2.2 s and after 8 s of UV exposure from paired recordings: *k*_des_ = 130 ± 15 s^−1^ (*n* = 9), *k*_des_ = 110 ± 10 s^−1^ (*n* = 9) and *k*_des_ = 50 ± 10 s^−1^ (*n* = 6); *I*_ss_/ *I*_peak_ = 0.03 ± 0.01 (n = 9), *I*_ss_/ *I*_peak_ = 0.05 ± 0.01 (n = 9) and *I*_ss_/ *I*_peak_ = 0.1 ± 0.02 (n = 6). **C** Exemplary normalized traces for responses to a 1 ms application of 10 mM glutamate, with the same color code as panel B. Bar graph shows rates of deactivation before, during and after UV, respectively: *k*_deac_ = 1400 ± 140 s^−1^ (*n* = 7), *k*_deac_ = 700 ± 140 s ^−1^ (*n* = 7) and *k*_deac_ = 500 ±140 s^−1^ (n = 4). Solid symbols indicate the rates for the responses shown in the left panels. Error bars represent SEM.

**Figure 9.**
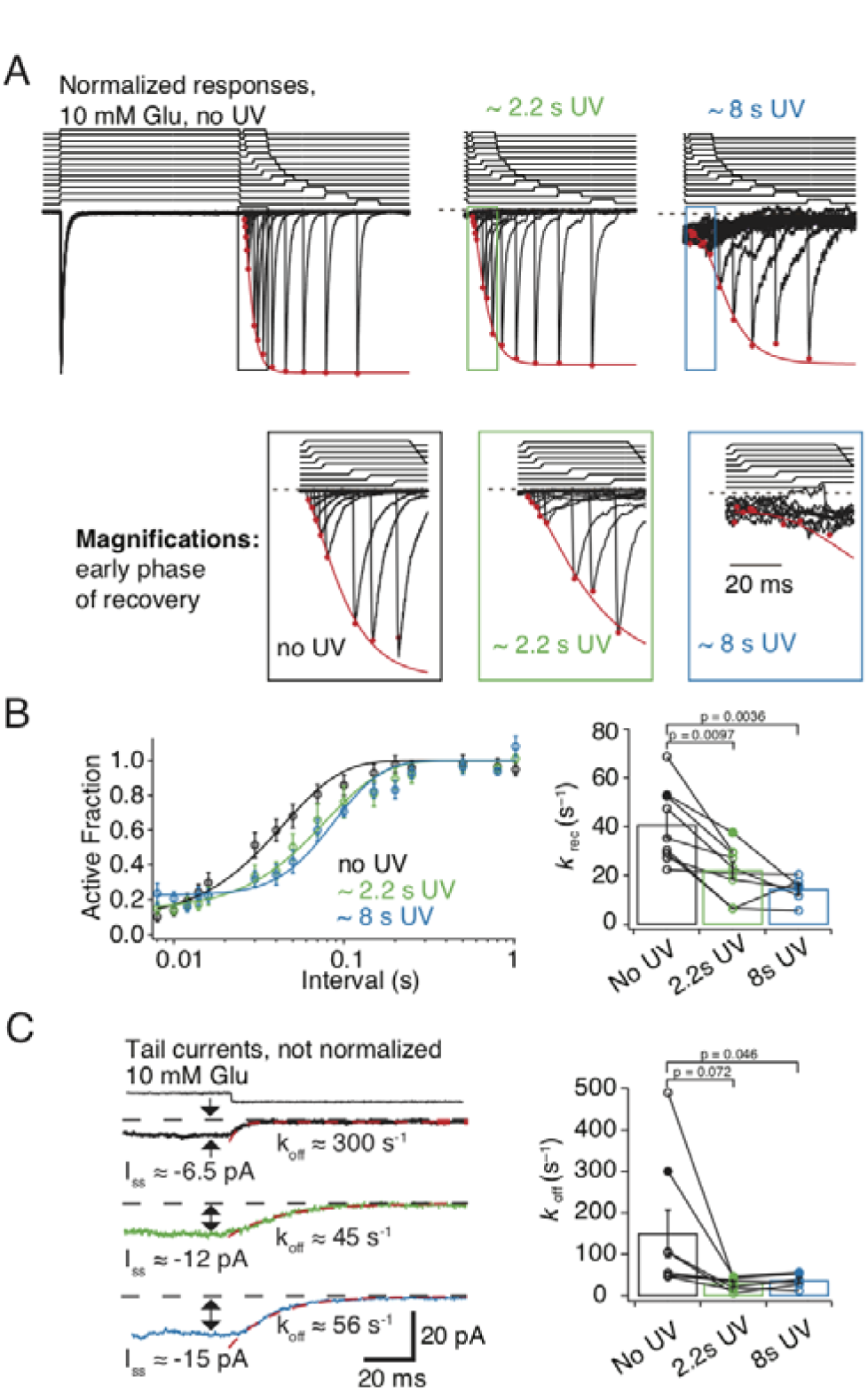
Role of the M2 segment in desensitized states. **A** Exemplary normalized traces for recovery from desensitization before, during (2.2 s, green) and after (8 s, blue) UV exposure. Time courses of glutamate applications are shown above the traces. Red circles indicate the peak of the response, fitted with a recovery function (red line). Apparent increase in noise following 8 s of UV exposure appears from performing a zoom on the trace, to be able to find the remaining glutamate induced current. **B** Recovery curves from pooled data are shifted to the right with cumulative UV exposure. Bar graph summarizing paired recovery from desensitization protocols before, during and after 8 s of UV exposure. Solid symbols refer to the patch in panel **A**. Recovery rates ± SEM *k*_rec_= 40 ± 5 s^−1^ (*n* = 9), 20 ± 3 s^−1^ (*n* = 9) and *k*_rec_ = 15 ± 2 s^−1^ (n = 6) before, during and after UV, respectively. **C** Exemplary traces for tail currents after desensitization. Bar graph of decays after the steady-state current before, during and after UV exposures, with solid symbols referring to the traces shown in the left panel. Deactivation rates ± SEM are *k*_off_ = 150 ± 60 s^−1^ (n = 9), *k*_off_ = 30 ± 5 s^−1^ (n=9) and *k*_off_ = 35 ± 10 s^−1^ (n = 6) before, during and after UV, respectively. Error bars represent SEM.

## Discussion

Despite the recent release of structures of activated GluA2 receptors (Twomey et al., 2017a; Chen et al., 2017) to compare with cognate closed state structures (Sobolevsky et al., 2009; Chen et al., 2014; Dϋrr et al., 2014), the nature and complexity of conformational changes occurring at the level of the TMD of AMPARs during gating remains unclear. Previous work supports the idea that M3 lines the pore and an upper hydrophobic box (Kuner et al., 2003; Alsaloum et al., 2016) and M2 comprises the selectivity filter region (Kuner et al., 2001). Conversely, the roles of the M1 and M4 helices are less well studied. In this work, site-specific incorporation of photoactivatable crosslinkers (Klippenstein et al., 2014) enabled access to the entire TMD of homomeric GluA2 AMPARs. A key advantage of this method is that function can be assessed before, during and after the photoactivation. Critically, crosslinking potential was independent of solvent exposure, allowing us to build a relatively unbiased map of functional elements in the TMD, covering sites that were previously inaccessible, and relate these elements to structural data. This factor was decisive in revealing complex relationships between the Pre-M1 and M4 helices and the selectivity filter region and channel activation and desensitization.

**Figure 10.**
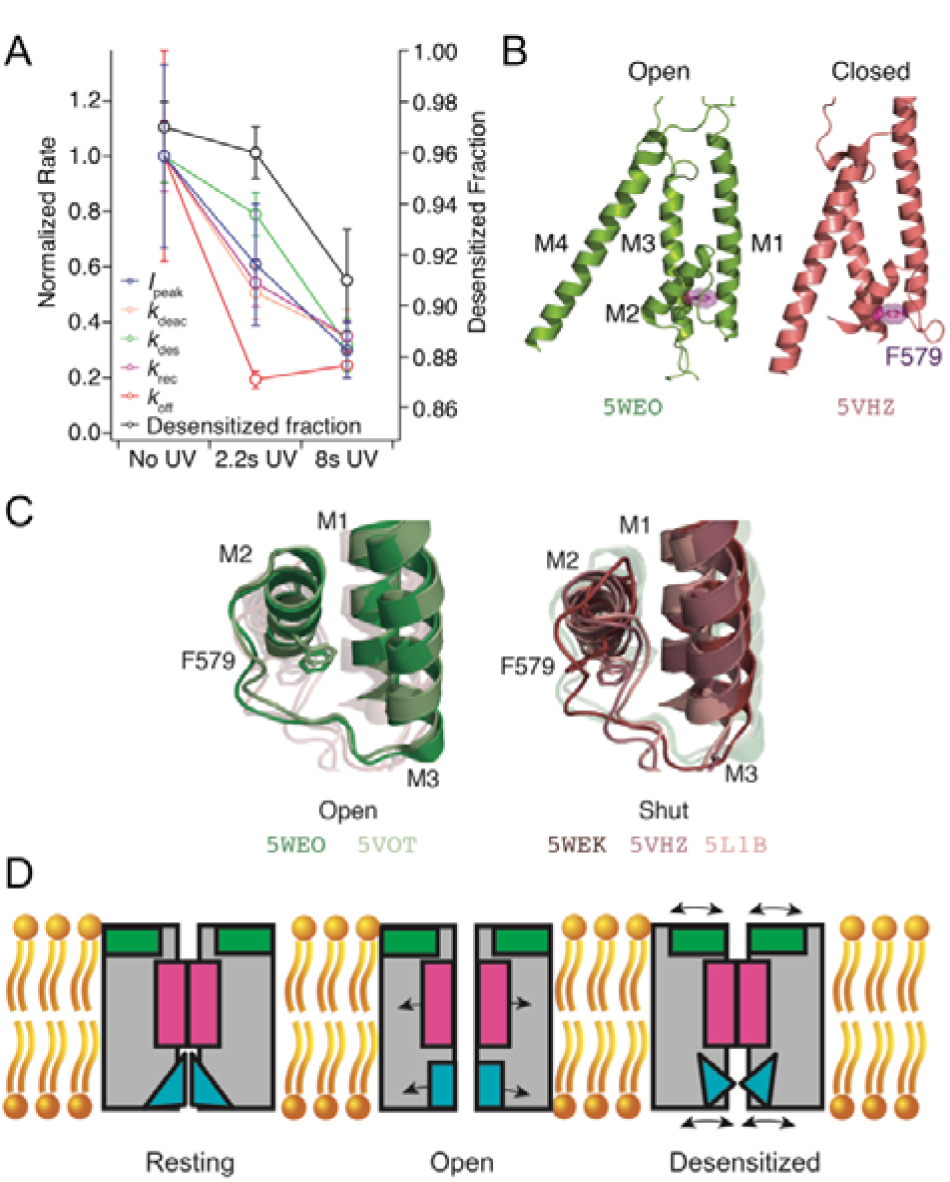
Putative AMPA receptor gating modules. **A** The time courses of UV-dependent gating changes in the F579AzF mutant (details in Figure 8 and 9). Desensitization changes (desensitized fraction and *k*_des_) developed over longer cumulative exposures than deactivation (*k*_deac_) or the long decay after a desensitized pulse (*k*_off_). For the inhibition of the peak current (*I*_peak_) before, during (2.2 s) and after (8s) UV exposure: *p* = 0.34 (paired t-test between no and 2.2s UV) and *p* = 0.0095 (repeated measures ANOVA). **B** The F579 site is located immediately behind the selectivity filter. Overlay of the TMD of a single subunit from closed (red) and open (green) GluA2 channel, with the four transmembrane helices (M1-4) indicated. **C** Overlays of open and closed channel structures showing F579 in multiple conformations. The PDB codes are indicated in the corresponding colors. **D** Scheme of gating modules. Channel opening requires a “bloom” at the M3 segment (gating module, magenta), and a conductive selectivity filter (cyan). Desensitization is accompanied by movements of the desensitization module (green) and potentially by structural dynamics of the selectivity filter.

The incorporation of bulky AzF and BzF residues had at most minor effects on the kinetics of channel activation, gating or desensitization of GluA2 before exposure to UV. The facility of these UAAs to be incorporated into the TMD depended on their environment and chemistry (Figure 1). Unsurprisingly, sites proximal to the phospholipid membrane (at the TMD periphery) were more likely to provide enough space for the insertion of both AzF and BzF, whereas the ion channel core was less permissive. However, the periphery was not insensitive to insertion of AzF and BzF, and I798AzF in M4 had, in contrast to other sites, basal kinetics (that is, before UV exposure) that differed from GluA2 WT. Following UV exposure, long lived (> 50 ms) single channel bursts could be observed for the I798AzF mutant (Supplementary Figure 8). These findings augment previous studies showing that insertion of tryptophan in sites of the peripheral TMD can disrupt transmembrane interactions and receptor tetramerization (Salussolia et al., 2011; Salussolia et al., 2013). At sites that were permissive, we found that approximately 50% of the AMPARs rescued by incorporation of AzF showed a change in channel gating upon UV exposure, while incorporation of BzF only showed an UV-induced effect in three cases out of 11 (Figure 1), two of which were at the membrane periphery. This superior capability of AzF in the TMD mig ht be due to its greater mobility to rotate and cross-link to nearby carbohydrogen, relative to BzF (Tian and Ye, 2016). Overall, UV-induced control of receptor activity was independent of the functional state and which membrane helix harboured the AzF residue. These observations favour the idea that all four membrane segments rearrange upon receptor activation in order to open the channel pore, or to permit receptor desensitization. The wide range of potential interacting partners for AzF or BzF in each state of activation may have precluded state-dependency. It is conceivable that UV illumination in different states result in crosslinking to various membrane segments or lipid parts of the bilayer, to yield the same effects.

Previous work suggested that a disulfide bond between the Pre-M1 and M4 helices could inhibit channel opening (Yelshanskaya et al., 2017). Four mutants clustered at the collar region (F515, L518, Y797, I798) had similar UV-dependent effects, unique to this gating module (Figure 7E, 10D). Our functional and biochemical data indicate that an adventitious crosslink, or interactions between Pre-M1 and M4, can also have a potentiation effect, and block desensitization (Figure 4). When BzF was incorporated in the likely solvent-exposed sites in the Pre-M1 helix (F515BzF and L518BzF) BzF crosslinking potentiated receptor currents (Figure 4). Biochemical analysis showed no intersubunit crosslinks, but we found a robust UV-induced increase in intrasubunit crosslinking, by quantifying monomeric protection of GluA2-F515BzF subunits cleaved by TEV protease. This result indicates a rotation of the Pre-M1 segment towards M4 of the same subunit (Figure 5D). This type of interaction has also been speculated for NMDA receptors (NMDARs) (Amin et al., 2017; Ogden et al., 2017). Furthermore, while NMDAR M4 segments have subunit-specific effects on desensitization, it has been shown that the extreme intracellular end of AMPAR M4 segments only contribute weakly to desensitization (Amin et al., 2017). Desensitization after photoactivation of Y797AzF was strongly reduced. Comparing this to our recent FRET data suggesting the M4 (as assayed from the C-terminal movement) is likely to move during gating, desensitization and without pore opening (Zachariassen et al., 2016), a wider role for peripheral interactions to modulate AMPAR function appears likely. We did not detect crosslinking either within or between subunits for Y797AzF. The effect in desensitization might thus be explained either by ring expansion of the AzF having a steric effect or by crosslinking to a lipid. We recently showed that interaction between the LBD-M4 linker and Stargazin is sufficient for modulation of GluA2 (Riva et al., 2017), thus raising the prospect that the perturbation at this site in M4 (Y797AzF), pointing away from the core of the channel, could be related to the changes in receptor function effected by auxiliary proteins, which in fact interact with the M4 and M1 of the AMPAR (Zhao et al., 2016; Twomey et al., 2017a). Notably, in some patches, we could detect that following photoactivation of AzF at position 798, glutamate could activate long bursts of channel openings with high open probability (Supplementary Figure 8), much like the incorporation of Stargazin produces (Tomita et al., 2005).

A distinct group of mutants flanking the bundle crossing (Figures 7E and 10D) showed properties likely resulting from an inhibition of the channel opening reaction, either by slowing channel opening, or destabilisation of the open state. F608 in the M3 helix showed the most potent inhibition. A kink at A618 allows the upper segment of M3 to move substantially upon channel opening (Twomey et al., 2017a), but F608 seems to move much less (Supplementary Figure 7). The fast rate of UV-inhibition at this site might therefore simply reflect the necessity of M3 movement to open/close the channel.

Both the structural and functional analyses of crosslinking sites indicated that F579AzF, located in close proximity to the selectivity filter at the base of the M2 helix, has a special role in gating of the receptor. In this context, the variation of the selectivity filter structure (and likewise its disordered nature in some other structures), including between closed state structures, and steric hindrance of the open channel selectivity filter structure by F579 are of interest (Figure 10C). Inhibition of the peak current was in the case of F579AzF accompanied by a large increase in the steady-state current. Mimicking these changes in a kinetic model required concurrent destabilization of both the open state and the desensitized state. The idea that desensitization can be affected by a residue deep in the channel in the M2 helix, was confirmed by the slowing of recovery from desensitization measured after UV exposure of GluA2-F579AzF (Figure 9). Notably, recovery happens entirely whilst the channel is closed. Therefore, crosslinking at this site had a dual effect on desensitization, slowing both entry and recovery (2.9-fold each), as well as separately biasing the open-closed equilibrium (because the peak current was substantially inhibited). Finally, we note that comparing our results to structures directly was generally unproductive (Supplementary Figure 6), whereas we readily discerned functional groups of residues by analyzing a 2-D plot of electrophysiological properties (Figure 7). This 2-D plot of peak vs. steady-state responses presumably gives insight because it compares the energies of modifying channel opening transitions and modifying receptor desensitization.

Overall, the effects of the F579AzF mutation following UV exposure are complex, and the extent of changes to the current have distinct dependencies on UV exposure. Our results do not distinguish between subunit dosage effects, that develop with a different dependence on the number of reacted subunits, or trapping of distinct structural forms. More work will be needed to dissect how the different UV-dependent chang es occur. The model that g ave the best description of the F579AzF data included progressive shift of the desensitized state towards a small conductance. In practice, such a phenomenon (the development of a new conductive state with UV exposure) may not be related to desensitization, but the relaxation from the steady-state current (from the tail current kinetics, Figure 9C) matched that of recovery from desensitization. However, our observations strongly corroborate the idea that the selectivity filter of AMPARs is as dynamic a structure as it is in simple tetrameric channels, with distinct arrangements between states (Twomey et al., 2017a). This property might allow the selectivity filter region to function as a second ion gate, closing and opening the pore between certain functional states as in other tetrameric channels. Further work will be required to assess whether state-dependent ion transit through the selectivity filter is a feature of the AMPAR transmembrane domain.

## Materials and Methods

### Molecular Biology

The aminoacyl-tRNA-synthetase and tRNA constructs for Human embryonic kidney (HEK) cell expression (Ye et al., 2008, Ye et al., 2009) were kind gifts from Thomas Sakmar (Rockefeller). For electrophysiological studies we used the pRK5 expression vector encoding the flip splice variant of the rat GluA2 subunit containing a Q at the Q/R-filter, followed by IRES and eGFP. The mutation Y40TAG was introduced into eGFP to act as a reporter of rescue (described in Klippenstein et al. 2014). Amber stop codons were introduced by overlap PCR and confirmed by double-stranded DNA sequencing. To study the active state of GluA2, a mutation (L483Y) was introduced that blocks receptor desensitization and stabilises an open state. (Sun et.al., 2002) The pRK5 vector was also used for biochemical experiments, but for this purpose the GluA2 subunit carried a C-terminal FLAG-tag epitope (Lau et al., 2013) for purification and three cysteines deleted (C190A, C436S, C528S) to lower background subunit dimerization (Klippenstein et al., 2014). The TEV protease recognition site ‘ENLYFQGS’ was inserted immediately before W572 in the M1-M2 intracellular loop with the native E571 being part of the TEV site (Xu et al., 2013).

### Cell culture and transfection

HEK293T for biochemical experiments and HEK293 cells for electrophysiological experiments were maintained in Miminum Essential Medium (MEM, Sigma-Aldrich) supplemented with 10% serum and 5% penicillin/streptomycin and grown at 37_°_C with 5% CO_2_. HEK293 and HEK293T cells were transiently transfected using polyethylenimine (PEI) in a 1:3 ratio (v/v; DNA/PEI) one day after cells were seeded. To suppress the amber stop codon, GluA2 mutants were co-expressed with vectors encoding mutated tRNA and synthetase for either AzF or BzF in the mass ratio 4:1:1. After six hours of incubation, the transfection medium was replaced by MEM supplemented with AzF (0.5 mM) or BzF (1 mM). We dissolved BzF (Bachem) in 1 M HCl and AzF (Chem-Impex International) in 1 M NaOH, which was immediately added to pre-warmed MEM containing 10% serum. Media supplemented with BzF or AzF were adjusted to pH 7.3 and filter-sterilized (0.22 µm PVDF filter) before use (Ye et al., 2009; Hino et al., 2006). Control experiments on wild-type receptors were done on the background of the AzF or BzF synthetase and the UAA medium.

### Electrophysiology

Patch clamp recordings of outside-out patches from HEK cells expressing mutant and wild-type glutamate receptors were performed 2-3 days after transfection. The external solution was composed as follows (mM): 150 NaCl, 0.1 MgCl_2_, 0.1 CaCl_2_, 5 HEPES. The pipette solution contained the following (mM): 115 NaCl, 0.5 CaCl_2_, 1 MgCl_2_, 10 Na_2_ATP, 10 NaF, 5 Na_4_BAPTA, 5 HEPES. Both solutions were titrated with NaOH to pH 7.3. Glutamate was diluted in the external solution and was applied to outside-out patches using a custom-made four-barrel glass perfusion tool (Vitrocom). For most experiments we used 10 mM glutamate to activate, but for some experiments, 1 mM glutamate was applied with no detectable differences in UV induced effects. The perfusion tool was mounted on a piezo-electric transducer (Physik Instrument), which was controlled via the digitizer interface (Instrutech ITC-18, HEKA Instrument). Borosilicate glass electrodes had resistance of 3-5 MΩ. Patches were clamped at −40 to −60 mV. Currents were filtered at 10 kHz and recorded at 40 kHz sampling rate using Axograph X (Axograph Scientific, Sydney, Australia). Macroscopic currents were elicited by applying the ligand for 400 ms. We exposed patches to UV light via epi-illumination from a Rapp UVICO source with a shutter under computer control. For state-dependent receptor trapping, patches were exposed to UV lig ht when the receptor was either in the resting (before g lutamate application), desensitised (during glutamate application) or in the active state (during glutamate application on L483Y-mutants) for 50 ms to 200 ms in each episode. It was not possible to record macroscopic currents from GluA2-F608AzF harbouring the L483Y mutation, so in this case 100 µM cyclothiazide (Ascent Scientific) was applied to trap the receptor in an active state. All chemicals were purchased from Roth unless otherwise noted.

### Kinetic Modelling

Simulated responses to glutamate for a set of kinetic models were generated using the Aligator scripts (www.github.com/aplested). Families of responses were generated by changing individual rate constants progressively, and correcting for microscopic reversibility on loops.These were adapted from previously used code to allow the conductance of states to change across a set of simulated currents, in order to mimic the desensitised state becoming conductive.

### Structural Analysis

We aligned the C-alpha atoms of residues 510-620 and 795-810 from chains A & C of GluA2 from six CryoEM and crystal structures: 5VOT, 5WEO (open), 5WEK, 5L1B (resting), 5VHZ, 5V0V (desensitised) (Yelshanskaya et al., 2016b; Twomey et al., 2017a; Twomey et al., 2017b; Chen et al., 2017) using the PyMol command “Align”. The selection of residues was interactively checked and chosen to give the most reliable alignment across all four subunits. Distances were taken between C-alpha atoms in the same chains for separate structures (for residue displacements), and between diametrically opposed chains in the same structures (for axial distances). Scripting was done in the Anaconda distribution of Python supplied with PyMol 2.0 (Schrödinger).

### Biochemistry

For biochemical experiments HEK293T cells were plated in dishes of 10 cm diameter and transfected as described above (cell culture and transfection). Three days after transfection the cells were exposed to UV light (~ 300 W/m^2^) on ice in a ventilated chamber (Luzchem, LZC-1) for 2-15 minutes. UV induced crosslinking was performed in the presence of 40 mM N-ethyl-maleimide (NEM, Thermo Fisher Scientific), to reduce spurious dimer formation from free cysteines following denaturation. Cells were harvested immediately after UV exposure, lysed in buffer containing 1% dodecylmaltoside (Glycon Biochemicals) and 40 mM NEM. Lysates were incubated with ANTI-FLAG M2 Affinity Gel (Sigma-Aldrich) and column purified (illustra MicroSpin, GE Healthcare). NEM inhibits TEV protease, therefore it was necessary to wash the samples extensively during purification to remove NEM before adding TEV protease for overnight digestion (TEV Protease, Protean). The second elution round was loaded on 4-12% NuPAGE Novex gels (Invitrogen) and run in reducing conditions in presence of 500 mM 2-Mercaptoethanol (Sigma-Aldrich) at 200 mV for 2 h. Transfer of the proteins to PVDF membranes (Immobilon-FL Millipore) was done using XCell surelock mini-cell and XCell II Blot Module (Invitrogen) as described in the manufacturer’s instructions. To enable quantitative detection of N - and C-terminal reactive bands in the same blot, membranes were incubated overnight at 4_°_C with polyclonal rabbit anti-GluR2/3 antibody (1:1000, Millipore) and monoclonal mouse anti-FLAG M2 antibody (1:1000, Sigma-Aldrich) respectively. The day after, infra-red dyes conjugated to secondary antibodies (IRDye 800CW Goat anti-mouse and IRDye 680RD Goat anti-Rabbit, Li-COR) were applied to the membranes for 1h at room temperature. The signal produced was detected on a LI-COR Odyssey Fc imager and quantified using ImageStudioLite2. As a control of readthrough the constructs containing an amber TAG stop codon at various positions in the GluA2 TMD were co-expressed with the tRNA and tRNA-synthetase in the absence of UAA, whereas WT controls were co-expressed with the tRNA and tRNA-synthetase in presence of UAA, to check for adventitious incorporation of UAA.

### Electrophysiology data analysis

Deactivation and desensitization were fit with either single or double exponential functions. To measure recovery from desensitization, two pulses of glutamate were applied in one episode with varying interpulse intervals. Recovery data were fit with a Hodgkin-Huxley type function (Carbone and Plested, 2012) with a slope of 2, except where noted. Kymograms of peak current reduction were fit with single exponential decay function. Statistical significance was assessed with Student’s t test, using either pairwise comparisons for different values from a single patch recording or unpaired tests for comparisons between mutants or different conditions. For multiple comparisons of the measurements before, during and after UV exposure of the F579AzF mutant, we performed repeated measures ANOVA.

## Acknowledgements

We are grateful to Thomas Sakmar (Rockefeller University) for providing the RNA-synthetase and tRNA plasmids. This work was funded by the Deutsche Forschungsgemeinschaft (DFG) Cluster of Excellence “NeuroCure” (EXC-257), DFG PL619-2 and SFB/TRR 186 to A.J.R.P. M.H.P. was recipient of fellowships from the Carlsberg foundation and the Danish Council for Independent Research. V.K. and A.P. both received stipends from the NeuroCure Cluster.

## Author contributions

All authors designed the experiments. All authors performed the experiments and/or analysed data. All authors read the manuscript. M.H.P., A.P. and A.J.R.P. wrote and edited the manuscript.

## Supplementary Material

### Movie 1 Gating modules

Residues corresponding to three spatially contiguous gating modules in the GluA2 pore domain are shown. The collar (green), the bundle crossing (magenta) and the selectivity filter (F579AzF, cyan) are flanked by the null mutants (wheat).

**Supplementary Table 1: Western blot analysis.**

**Supplementary Table 2: Kinetics of glutamate responses of selected constructs.**

**Supplementary Table 3: Summary of UV-induced changes in glutamate response.**

**Supplementary Table 4: Amplitudes of control and “readthrough” currents.**

**Supplementary Figure 1.**
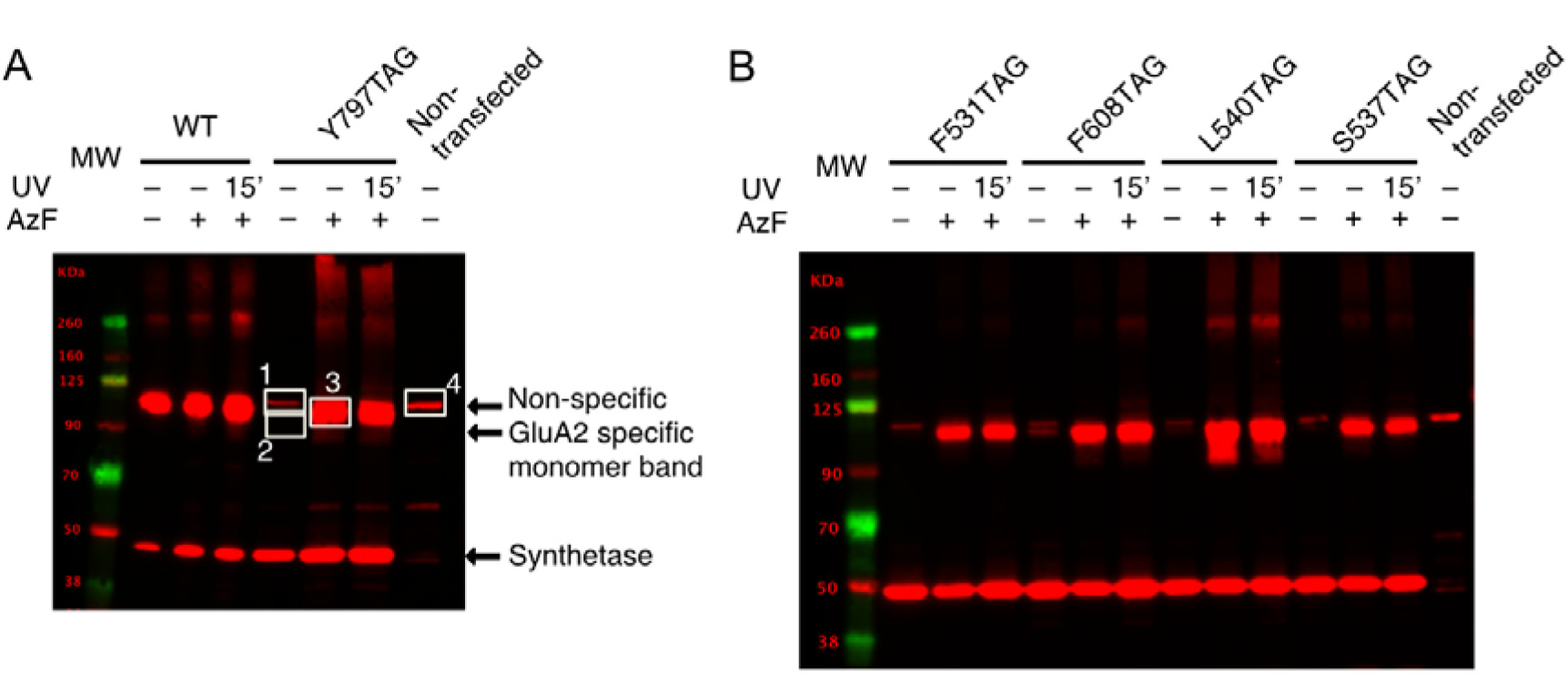
Representative Western blots of GluA2 TAG mutants rescued by AzF incorporation. **A** Estimation of the enrichment of expression by AzF for the Y797TAG mutant. Western blotting against the FLAG epitopes on GluA2 and the tRNA synthetase also reveals a non-specific band around 100 kDa, similar in size to monomeric GluA2 (indicated for non-transfected HEK 293-T cells with white box #4). Since we could not separate the non-specific band from actual rescue in presence of UAA, we calculated minimum enrichment following inclusion of AzF by taking the ratio of densitometric measurement including the non-specific band (Box #3 / [Box #1 + Box #2]) or the maximum enrichment by excluding the non-specific band (Box #3/ Box #2). **B** Typical Western blot for measurement of enrichment following AzF inclusion in culture media for four TAG mutants of GluA2. Dimeric fractions can be seen around 260 kDa for L540, but these were not enhanced by UV (15’ exposure).

**Supplementary Figure 2.**
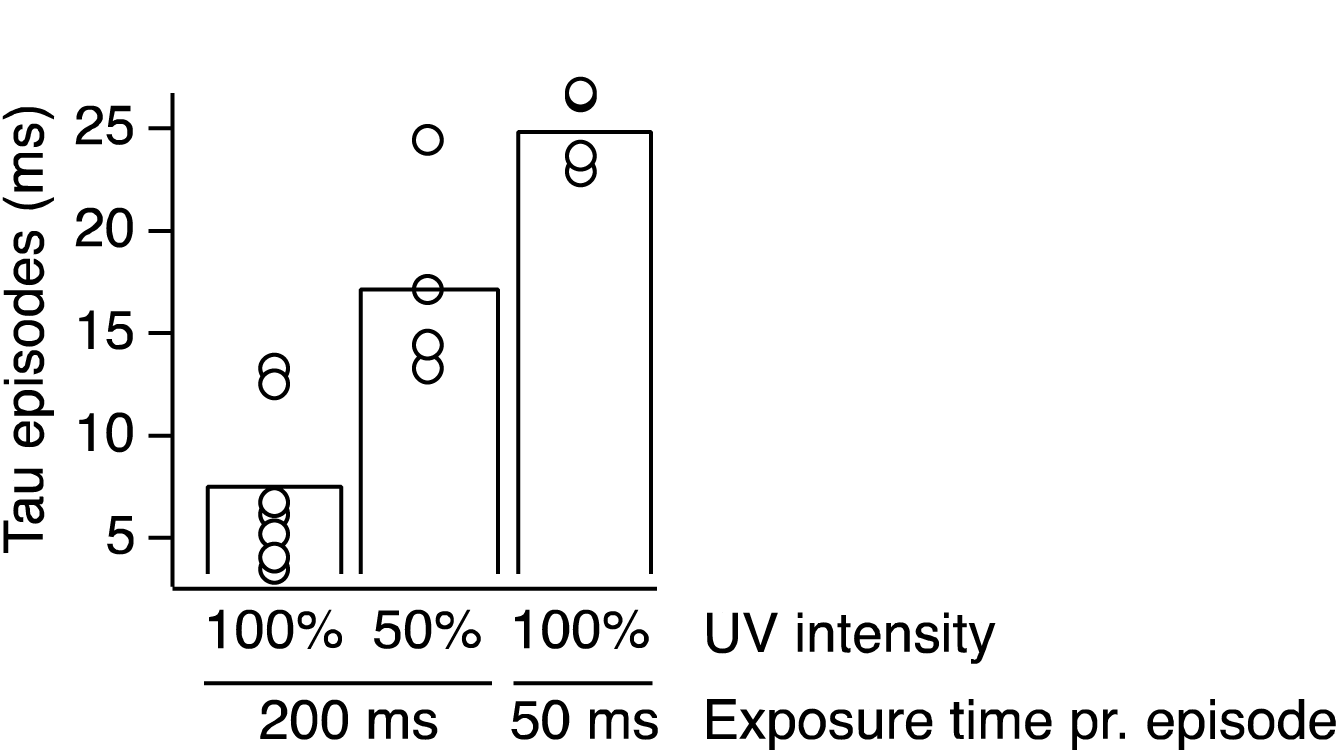
Dependency on UV intensity and exposure time. Summary of the exponential half-times of GluA2-F608AzF inactivation, plotted against the UV exposure periods per episode in milliseconds. The rate of peak current reduction could be manipulated by changing the intensity of the UV light from 100% (τ 200ms,100% = 7 ms) to 50% (τ 200ms,50% = 17 ms) or reducing the time interval of UV exposure to 50 ms (τ 50ms,100% = 25 ms), verifying that the peak-current reduction is controlled by UV light.

**Supplementary Figure 3.**
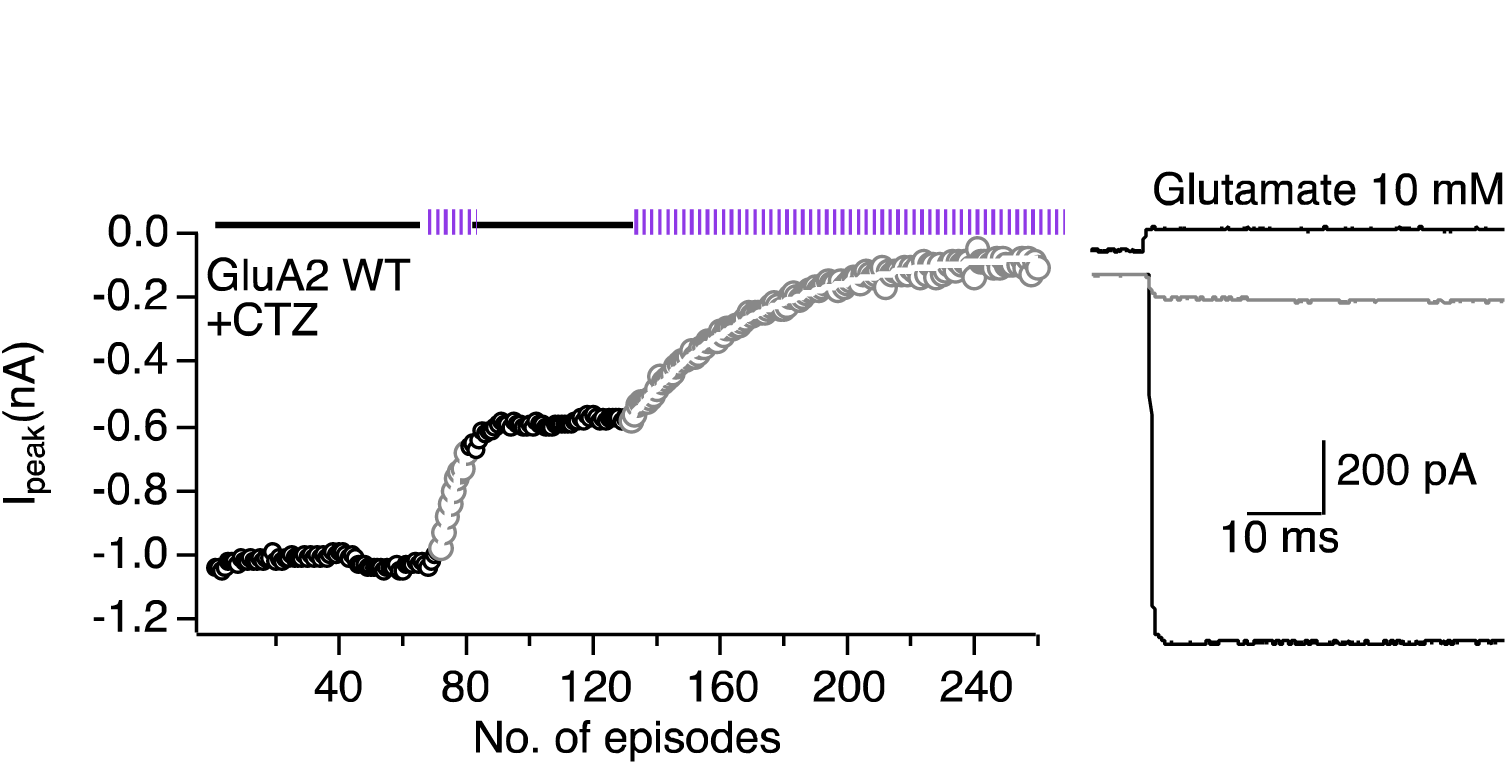
UV-induced inhibition of GluA2 wild-type in the presence of CTZ. UV exposures resulted in a decrease in the peak current after the application of glutamate and 100 µM CTZ. This effect was irregular and showed batch to batch variation (compare with Klippenstein et al. 2014). *Left*; Example kymogram illustrates the UV inactivation of GluA2 WT due to the presence of CTZ, plotted as described in the legend to **Figure 3**. *Right*; Example responses from the beginning (black) and end of the kymogram (grey).

**Supplementary Figure 4.**
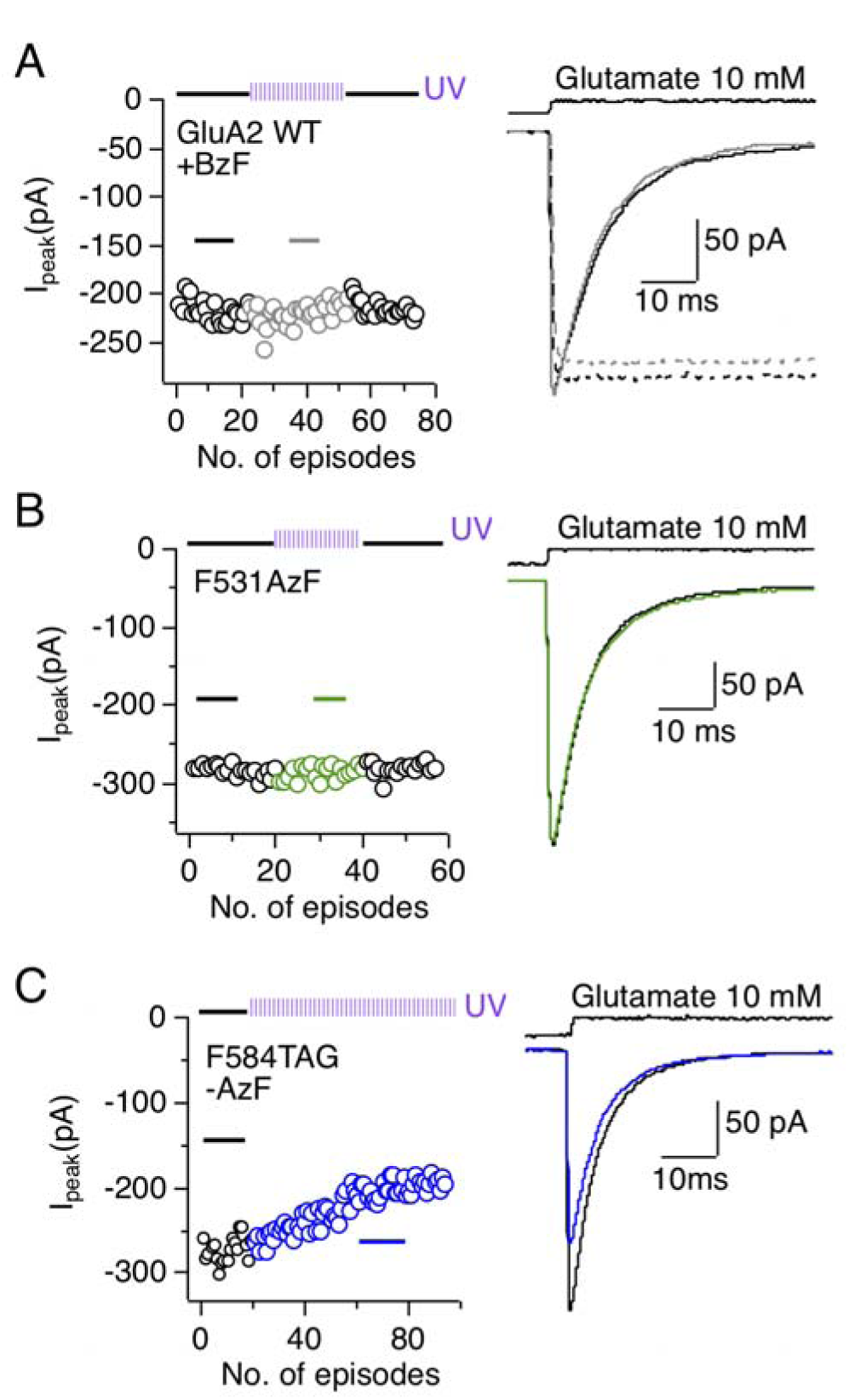
Three null results of UV exposure. **A** UV exposure did not change the glutamate response for the GluA2 WT cultured in the presence of BzF and the appropriate tRNA-synthetase pair. Kymogram (*left*) as described in Figure 3A-E illustrates the time course of UV exposure, and example currents (*right*) from sections of the histogram labeled with a solid line. The effect of UV exposure in the active state of the GluA2 WT was also tested by introducing the L483Y mutation to block desensitization (dotted lines). **B** GluA2-F531AzF was insensitive to UV. **C** All tested TAG mutants that were not rescued with an UAA, but that gave a current due to read through, were also insensitive to UV, as exemplified by F584TAG. This patch exhibited a constant mild rundown, independent of the UV exposure.

**Supplementary Figure 5.**
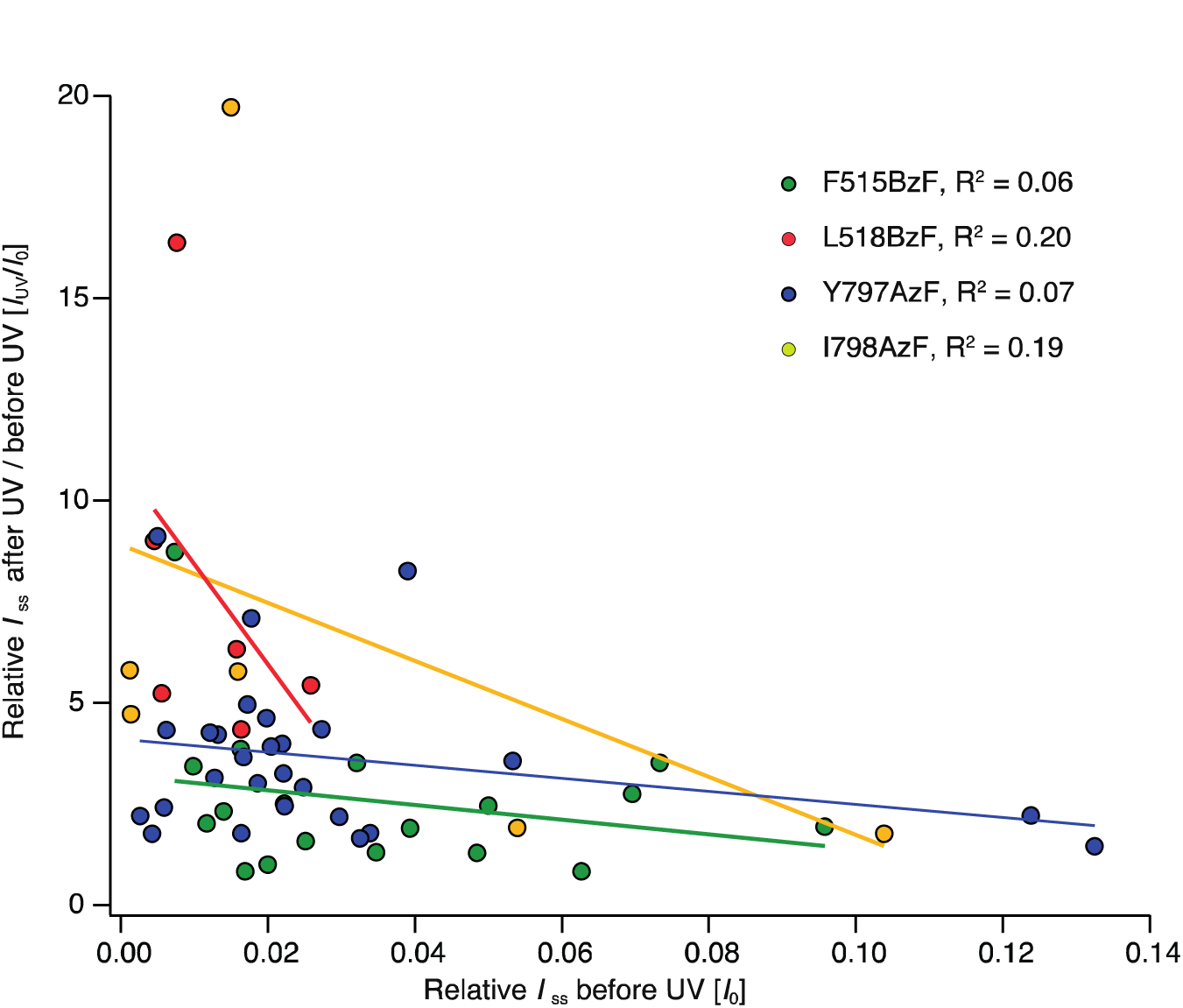
Relation between the initial steady-state current and phtotopotentiation. Graph showing the absence of correlation between relative steady-state currents (I_ss_/I_peak_) before UV application (I_0_) and the fold change in relative steady-state current (Iss/Ipeak) before and after UV application (I_UV_/I_0_) for the GluA2-F515BzF, - L518BzF, - Y797AzF and - I798AzF mutants. Thus, UV induced photopotentiation of the relative steady-state current was independent of the amplitude of the initial relative steady-state current.

**Supplementary Figure 6.**
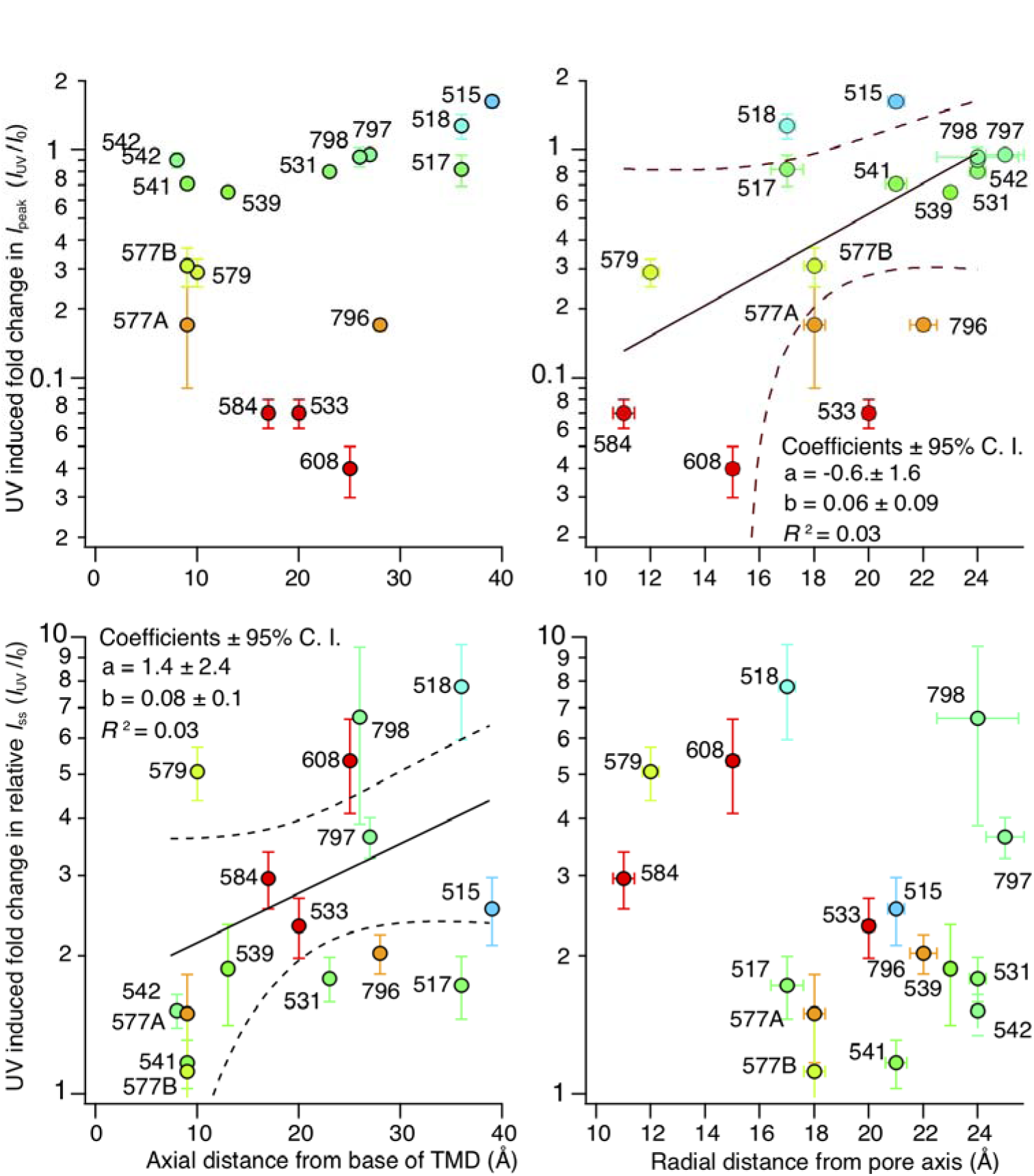
Comparing distances at photoactive sites and UV induced effects. The UV induced fold changes of peak currents (*upper panels*)and relative steady-state currents (I_ss_/I_peak_) (*lower panels*), of each mutant were plotted against the axial distance of UAA insertion site from the base of the TMD (*left panels*) or the radial distance from the pore axis (*right panels*) in Ångström. Distances were taken between the C_α_ atoms of the UAA insertion site and the base of the TMD (L817) or between C_α_ atoms of the UAA insertion site in diametrically opposed chains. The sites are color-coded according to their UV-induced fold change in peak current (as in Figure 6B). 577A and 577B denotes the insertion of AzF and BzF, respectively, at this particular site.

**Supplementary Figure 7.**
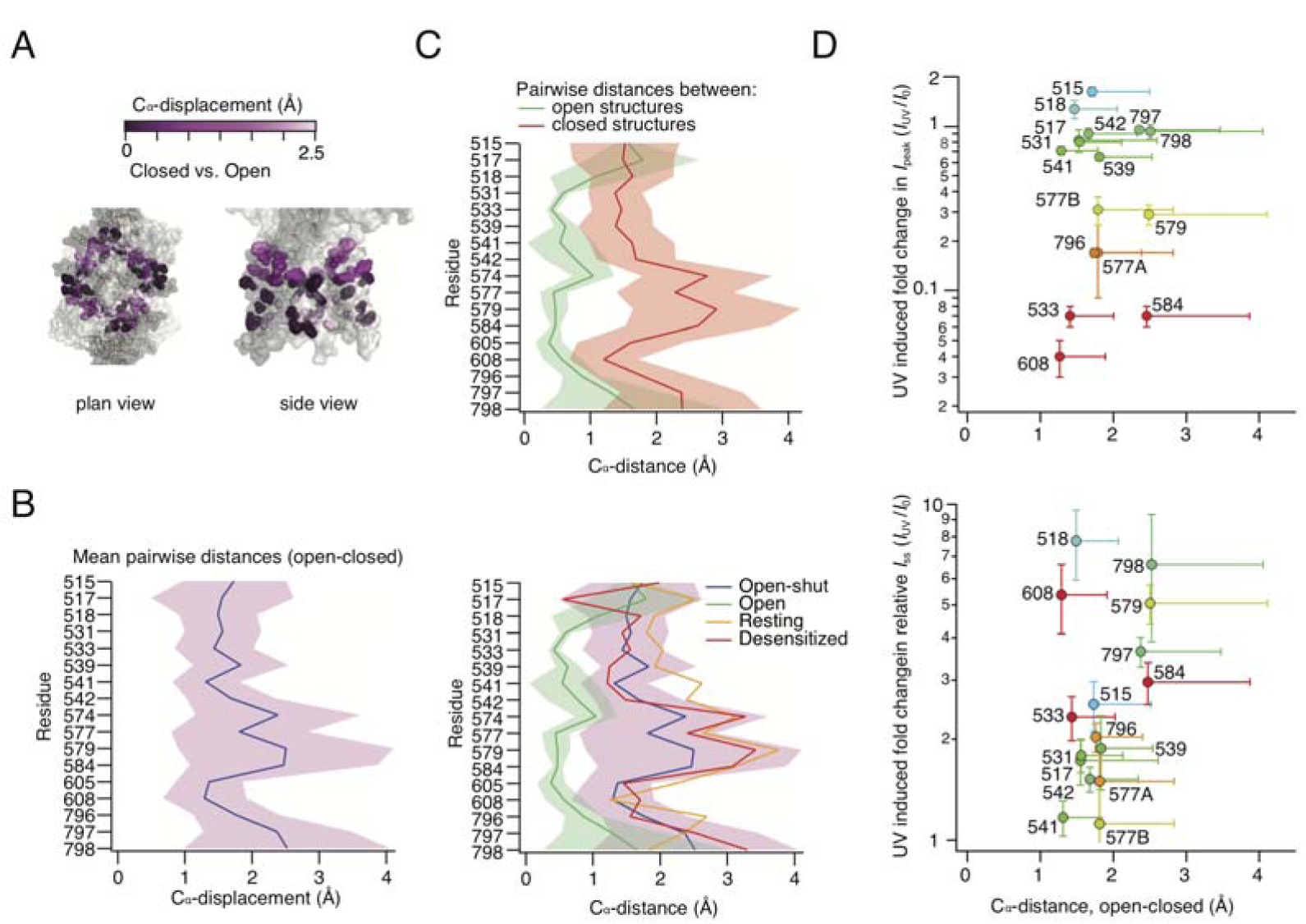
Amino acid displacements in published structures. **A** Colored bar (*upper*) shows the used color code for C_α_-displacement in Ångström of the AzF/BzF insertion sites between resting (PDB ID: 3kg2) and active (PDB ID: 5weo) state. Dark magenta indicates no to very small movements, whereas light magenta indicates bigger movements. Pymol figures (*bottom*) show selected insertion sites as spheres colored according to its UV C_α_-displacement effect. **B** Relation between the C_α_;-distances at the UAA insertion sites in the open to closed state structures. **C** Structures were further divided into closed and open (*upper panel*) and open-shut, open, resting and desensitized state (*lower panel*). **D** Graph showing the C _α_-distance between open and closed structures and the site-specific UV-induced effect on receptor peak current amplitude (*upper*) and relative steady-state current (*bottom*) for each site of the TMD, respectively. The individual sites are color-coded according to the specific UV-induced effects on peak current as in supplementary figure 6. Distances between C_α_atoms in the same chains were measured between open and closed channel structures. Errors derive from variability in distances between different subunits and between different structures.

**Supplementary Figure 8.**
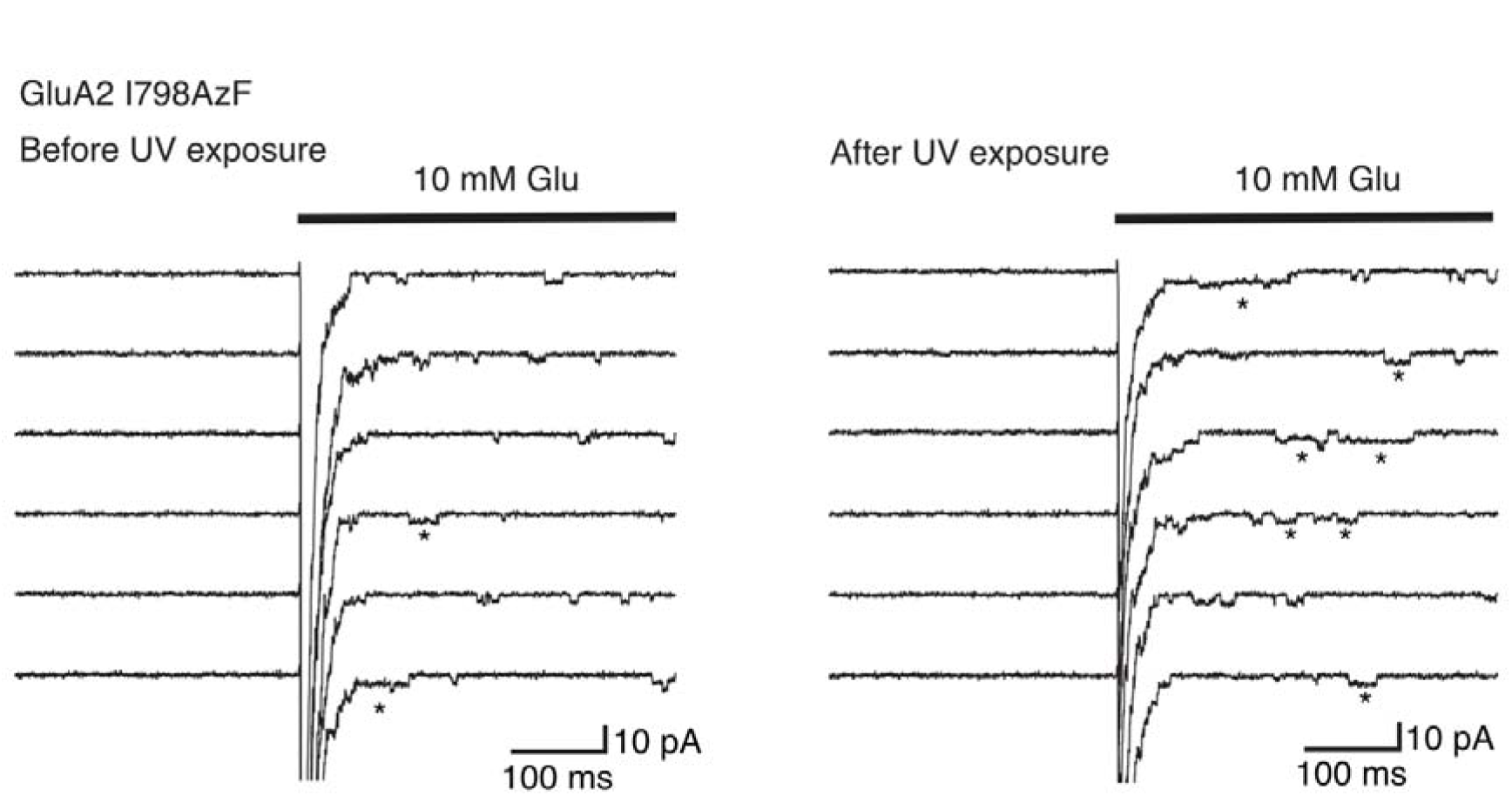
Photoactivation at I798AzF lengthens the activations of individual receptors. Representative desensitized traces from GluA2-I798AzF outside-out patches before (*left*) and after (*right*) UV exposure. After UV exposure, long bursts (>25 ms) with high open probability (*) became much more prevalent. The traces were filtered at 1kHz. The holding voltage was −60 mV.

## Bibliography

Ahmed, A. H., Wang, S., Chuang, H.-H., and Oswald, R. E. (2011). Mechanism of AMPA receptor activation by partial agonists: disulfide trapping of closed lobe conformations. J Biol Chem 286, 35257–35266.

Alsaloum, M., Kazi, R., Gan, Q., Amin, J., and Wollmuth, L. P. (2016). A Molecular Determinant of Subtype-Specific Desensitization in Ionotropic Glutamate Receptors. J Neurosci 36, 2617–2622.

Amin, J. B., Salussolia, C. L., Chan, K., Regan, M. C., Dai, J., Zhou, H. X., Furukawa, H., Bowen, M. E., and Wollmuth, L. P. (2017). Divergent roles of a peripheral transmembrane segment in AMPA and NMDA receptors. J Gen Physiol 149, 661–680.

Armstrong, N., Jasti, J., Beich-Frandsen, M., and Gouaux, E. (2006). Measurement of conformational changes accompanying desensitization in an ionotropic glutamate receptor. Cell 127, 85–97.

Baranovic, J., Chebli, M., Salazar, H., Carbone, A. L., Faelber, K., Lau, A. Y., Daumke, O., and Plested, A. J. (2016). Dynamics of the Ligand Binding Domain Layer during AMPA Receptor Activation. Biophys J 110, 896–911.

Blunck, R., Cordero-Morales, J. F., Cuello, L. G., Perozo, E., and Bezanilla, F. (2006). Detection of the opening of the bundle crossing in KcsA with fluorescence lifetime spectroscopy reveals the existence of two gates for ion conduction. J Gen Physiol 128, 569–581.

Carbone, A. L., and Plested, A. J. R. (2012). Coupled control of desensitization and gating by the ligand binding domain of glutamate receptors. Neuron 74, 845–857.

Carbone, A. L., and Plested, A. J. R. (2016). Superactivation of AMPA receptors by auxiliary proteins. Nat Commun 7, 10178.

Chen, L., Dürr, K. L., and Gouaux, E. (2014). X-ray structures of AMPA receptor-cone snail toxin complexes illuminate activation mechanism. Science 345, 1021–1026.

Chen, S., Zhao, Y., Wang, Y., Shekhar, M., Tajkhorshid, E., and Gouaux, E. (2017). Activation and Desensitization Mechanism of AMPA Receptor-TARP Complex by Cryo-EM. Cell 170, 1234–1246.e14.

Contreras, J. E., Srikumar, D., and Holmgren, M. (2008). Gating at the selectivity filter in cyclic nucleotide-gated channels. Proceedings of the National Academy of Sciences 105, 3310.

del Camino, D., and Yellen, G. (2001). Tight Steric Closure at the Intracellular Activation Gate of a Voltage-Gated K+ Channel. Neuron 32, 649–656.

Devaraneni, P. K., Komarov, A. G., Costantino, C. A., Devereaux, J. J., Matulef, K., and Valiyaveetil, F. I. (2013). Semisynthetic K+ channels show that the constricted conformation of the selectivity filter is not the C-type inactivated state. Proc Natl Acad Sci U S A 110, 15698–15703.

Dϋrr, K. L., Chen, L., Stein, R. A., De Zorzi, R., Folea, I. M., Walz, T., Mchaourab, H. S., and Gouaux, E. (2014). Structure and Dynamics of AMPA Receptor GluA2 in Resting, Pre-Open, and Desensitized States. Cell 158, 778–792.

Grosman, C., Zhou, M., and Auerbach, A. (2000). Mapping the conformational wave of acetylcholine receptor channel gating. Nature 403, 773–776.

Hino, N., Hayashi, A., Sakamoto, K., and Yokoyama, S. (2006). Site-specific incorporation of non-natural amino acids into proteins in mammalian cells with an expanded genetic code. Nat Protoc 1, 2957–2962.

Horning, M. S., and Mayer, M. L. (2004). Regulation of AMPA receptor gating by ligand binding core dimers. Neuron 41, 379–388.

Klippenstein, V., Hoppmann, C., Ye, S., Wang, L., and Paoletti, P. (2017). Optocontrol of glutamate receptor activity by single side-chain photoisomerization. Elife 6,

Klippenstein, V., Ghisi, V., Wietstruk, M., and Plested, A. J. R. (2014). Photoinactivation of glutamate receptors by genetically encoded unnatural amino acids. J Neurosci 34, 980–991.

Kuner, T., Beck, C., Sakmann, B., and Seeburg, P. H. (2001). Channel-lining residues of the AMPA receptor M2 segment: structural environment of the Q/R site and identification of the selectivity filter. J Neurosci 21, 4162–4172.

Kuner, T., Seeburg, P. H., and Guy, H. R. (2003). A common architecture for K+ channels and ionotropic glutamate receptors. Trends Neurosci 26, 27–32.

Labro, A. J., Cortes, D. M., Tilegenova, C., and Cuello, L. G. (2018). Inverted allosteric coupling between activation and inactivation gates in K_+_ channels. Proc Natl Acad Sci U S A 115, 5426–5431.

Lau, A. Y., Salazar, H., Blachowicz, L., Ghisi, V., Plested, A. J. R., and Roux, B. (2013). A conformational intermediate in glutamate receptor activation. Neuron 79, 492–503.

Martin, G. M., Rex, E. A., Devaraneni, P., Denton, J. S., Boodhansingh, K. E., DeLeon, D. D., Stanley, C. A., and Shyng, S. L. (2016). Pharmacological Correction of Trafficking Defects in ATP-sensitive Potassium Channels Caused by Sulfonylurea Receptor 1 Mutations. J Biol Chem 291, 21971–21983.

Murray, C. I., Westhoff, M., Eldstrom, J., Thompson, E., Emes, R., and Fedida, D. (2016). Unnatural amino acid photo-crosslinking of the IKs channel complex demonstrates a KCNE1:KCNQ1 stoichiometry of up to 4:4. Elife 5,

Naganathan, S., Ye, S., Sakmar, T. P., and Huber, T. (2013). Site-specific epitope tagging of G protein-coupled receptors by bioorthogonal modification of a genetically encoded unnatural amino acid. Biochemistry 52, 1028–1036.

Oelstrom, K., Goldschen-Ohm, M. P., Holmgren, M., and Chanda, B. (2014). Evolutionarily conserved intracellular gate of voltage-dependent sodium channels. Nat Commun 5, 3420.

Ogden, K. K., Chen, W., Swanger, S. A., McDaniel, M. J., Fan, L. Z., Hu, C., Tankovic, A., Kusumoto, H., Kosobucki, G. J., Schulien, A. J., Su, Z., Pecha, J., Bhattacharya, S., Petrovski, S., Cohen, A. E., Aizenman, E., Traynelis, S. F., and Yuan, H. (2017). Molecular Mechanism of Disease-Associated Mutations in the Pre-M1 Helix of NMDA Receptors and Potential Rescue Pharmacology. PLoS Genet 13, e1006536.

Posson, D. J., McCoy, J. G., and Nimigean, C. M. (2013). The voltage-dependent gate in MthK potassium channels is located at the selectivity filter. Nat Struct Mol Biol 20, 159–166.

Riva, I., Eibl, C., Volkmer, R., Carbone, A. L., and Plested, A. J. (2017). Control of AMPA receptor activity by the extracellular loops of auxiliary proteins. Elife 6,

Salussolia, C. L., Corrales, A., Talukder, I., Kazi, R., Akgul, G., Bowen, M., and Wollmuth, L. P. (2011). Interaction of the M4 segment with other transmembrane segments is required for surface expression of mammalian α-amino-3-hydroxy-5-methyl-4-isoxazolepropionic acid (AMPA) receptors. J Biol Chem 286, 40205–40218.

Salussolia, C. L., Gan, Q., Kazi, R., Singh, P., Allopenna, J., Furukawa, H., and Wollmuth, L. P. (2013). A eukaryotic specific transmembrane segment is required for tetramerization in AMPA receptors. J Neurosci 33, 9840–9845.

Sobolevsky, A. I., Yelshansky, M. V., and Wollmuth, L. P. (2004). The outer pore of the glutamate receptor channel has 2-fold rotational symmetry. Neuron 41, 367–378.

Sobolevsky, A. I., Rosconi, M. P., and Gouaux, E. (2009). X-ray structure, symmetry and mechanism of an AMPA-subtype glutamate receptor. Nature 462, 745–756.

Sobolevsky, A. I., Yelshansky, M. V., and Wollmuth, L. P. (2003). Different gating mechanisms in glutamate receptor and K+ channels. J Neurosci 23, 7559–7568.

Sobolevsky, A. I., Yelshansky, M. V., and Wollmuth, L. P. (2005). State-dependent changes in the electrostatic potential in the pore of a GluR channel. Biophys J 88, 235–242.

Sun, Y., Olson, R., Horning, M., Armstrong, N., Mayer, M., and Gouaux, E. (2002). Mechanism of glutamate receptor desensitization. Nature 417, 245–253.

Thompson, J., and Begenisich, T. (2012). Selectivity filter gating in large-conductance Ca(2+)-activated K+ channels. J Gen Physiol 139, 235–244.

Tian, M., and Ye, S. (2016). Allosteric regulation in NMDA receptors revealed by the genetically encoded photo-cross-linkers. Sci Rep 6, 34751.

Tilegenova, C., Cortes, D. M., and Cuello, L. G. (2017). Hysteresis of KcsA potassium channel’s activation - deactivation gating is caused by structural changes at the channel’s selectivity filter. Proc Natl Acad Sci U S A 114, 3234–3239.

Tomita, S., Adesnik, H., Sekiguchi, M., Zhang, W., Wada, K., Howe, J. R., Nicoll, R. A., and Bredt, D. S. (2005). Stargazin modulates AMPA receptor gating and trafficking by distinct domains. Nature 435, 1052–1058.

Twomey, E. C., Yelshanskaya, M. V., Grassucci, R. A., Frank, J., and Sobolevsky, A. I. (2017a). Channel opening and gating mechanism in AMPA-subtype glutamate receptors.

Nature Twomey, E. C., Yelshanskaya, M. V., Grassucci, R. A., Frank, J., and Sobolevsky, A. I. (2017b). Structural Bases of Desensitization in AMPA Receptor-Auxiliary Subunit Complexes. Neuron 94, 569–580.e5.

Xu, Y., Ramu, Y., Shin, H.-G., Yamakaze, J., and Lu, Z. (2013). Energetic role of the paddle motif in voltage gating of Shaker K(+) channels. Nat Struct Mol Biol 20, 574–581.

Ye, S., Huber, T., Vogel, R., and Sakmar, T. P. (2009). FTIR analysis of GPCR activation using azido probes. Nat Chem Biol 5, 397–399.

Ye, S., Köhrer, C., Huber, T., Kazmi, M., Sachdev, P., Yan, E. C. Y., Bhagat, A., RajBhandary, U. L., and Sakmar, T. P. (2008). Site-specific incorporation of keto amino acids into functional G protein-coupled receptors using unnatural amino acid mutagenesis. J Biol Chem 283, 1525–1533.

Ye, S., Zaitseva, E., Caltabiano, G., Schertler, G. F. X., Sakmar, T. P., Deupi, X., and Vogel, R. (2010). Tracking G-protein-coupled receptor activation using genetically encoded infrared probes. Nature 464, 1386–1389.

Yelshanskaya, M. V., Saotome, K., Singh, A. K., and Sobolevsky, A. I. (2016a). Probing Intersubunit Interfaces in AMPA-subtype Ionotropic Glutamate Receptors. Sci Rep 6, 19082.

Yelshanskaya, M. V., Mesbahi-Vasey, S., Kurnikova, M. G., and Sobolevsky, A. I. (2017). Role of the Ion Channel Extracellular Collar in AMPA Receptor Gating. Sci Rep 7, 1050.

Yelshanskaya, M. V., Singh, A. K., Sampson, J. M., Narangoda, C., Kurnikova, M., and Sobolevsky, A. I. (2016b). Structural Bases of Noncompetitive Inhibition of AMPA-Subtype Ionotropic Glutamate Receptors by Antiepileptic Drugs. Neuron 91, 1305–1315.

Zachariassen, L. G., Katchan, L., Jensen, A. G., Pickering, D. S., Plested, A. J., and Kristensen, A. S. (2016). Structural rearrangement of the intracellular domains during AMPA receptor activation. Proc Natl Acad Sci U S A 113, E3950–9.

Zhao, Y., Chen, S., Yoshioka, C., Baconguis, I., and Gouaux, E. (2016). Architecture of fully occupied GluA2 AMPA receptor-TARP complex elucidated by cryo-EM.

Nature Zhou, Y., Xia, X. M., and Lingle, C. J. (2011). Cysteine scanning and modification reveal major differences between BK channels and Kv channels in the inner pore region. Proc Natl Acad Sci U S A 108, 12161–12166.

